# Structural Insights into Regulation of Insulin Expression Involving i-Motif DNA Structures in the Insulin-Linked Polymorphic Region

**DOI:** 10.1101/2023.06.01.543149

**Authors:** Dilek Guneri, Effrosyni Alexandrou, Kamel El Omari, Zuzana Dvořáková, Rupesh V. Chikhale, Daniel Pike, Christopher A. Waudby, Christopher J. Morris, Shozeb Haider, Gary N. Parkinson, Zoë A. E. Waller

## Abstract

The insulin linked polymorphic region (ILPR) is a variable number of tandem repeats (VNTR) region of DNA in the promoter of the insulin gene that regulates transcription of insulin. This region is known to form the alternative DNA structures, i-motifs and G-quadruplexes. Individuals have different sequence variants of VNTR repeats and although previous work investigated the effects of some variants on G-quadruplex formation, there is not a clear picture of the relationship between the sequence diversity, the DNA structures formed, and the functional effects on insulin gene expression. Here we show that different sequence variants of the ILPR form different DNA secondary structures and insulin expression is dependent on formation of i-motif and G-quadruplex structures. The first crystal structure and dynamics of an intramolecular i-motif also reveal sequences within the loop regions forming additional stabilising interactions, which are critical to formation of the stable i-motif structures that modulate insulin expression. The outcomes of this work reveal the detail in formation of stable i-motif DNA structures, with potential for rational based drug design for compounds to alter insulin gene expression.

## INTRODUCTION

Insulin (INS) is a protein hormone central to the regulation of glucose metabolism. Deficiencies or incorrect production of insulin can lead to hyperglycemia and diabetes mellitus.^1, 2^ The insulin linked polymorphic region (ILPR) is a variable number of tandem repeats (VNTR) region of DNA in the promoter of the insulin gene that regulates transcription of insulin.^3, 4^ The ILPR sequence is a minisatellite located 363 bp upstream of the insulin transcription start site with heterogeneity in the number of tandemly repeated sequences observed among individuals.^4^ The predominant ILPR sequence is composed of 14 base pair tandem repeats comprising of 5’-ACAGGGGTGTGGGG-3’/3’-TGTCCCCACACCCC-5’.^4^ Shortening in the length of the ILPR and variants in the sequence has been linked to development of both Type-1 and Type-2 diabetes.^5–9^ The ILPR influences both the expression of INS and insulin-like growth factor 2 and genetic variations in the ILPR are also associated with decreased expression of INS.^5, 10, 11^ It has also been shown that insulin itself binds the G-rich regions in the ILPR.^12^ However, exactly how the changes in the ILPR cause alterations in expression of the INS gene remains unclear.

The ILPR is a GC-rich region of DNA.^3–5, 13, 14^ The C-rich sequence can form i-motif structures, comprised of two parallel stranded hairpins zipped together by intercalated, hemiprotonated cytosine-cytosine base pairs.^15^ In contrast, the G-rich sequence can fold into G-quadruplexes, which form from planar G-quartets, held together by Hoogsteen hydrogen bonding and further stabilised by via π–π stacking and the coordination of cations.^16^ These types of non-canonical secondary structures have been shown to exist in cells and are prevalent within the promoter regions of genes, in particular, regions linked to regulation of gene expression and other regulatory elements.^17, 18^ Although G-quadruplex structures have been well studied, much less research has focussed on i-motifs, the sequences that comprise them, the corresponding structures they form and their biological functions.

Herein we characterise the different variants of the C-rich and G-rich sequences within the ILPR and determine a relationship between the stable formation of i-motif and G-quadruplex structures with corresponding insulin gene expression. The first crystal structure of an intramolecular i-motif also reveals that the sequences within the loop regions, and their additional stabilising interactions, are critical to formation of the stable i-motif structures that control insulin expression.

## RESULTS AND DISCUSSION

### Characterisation of the Sequence Variants Within the ILPR

The polymorphism of the ILPR sequence extends to at least 11 main variations, with minor changes in the loops or C/G-tracts from the predominant ILPR sequence.^15^ Some of these variations were shown to express different levels of insulin compared to the most prevalent ILPR sequence.^5^ Some previous work had studied three of the most prevalent G-rich variants of these sequences and were able to correlate a relationship between the conformation of the G-quadruplex structure with binding affinity to insulin and insulin-like growth factor.^19^ However, to fully understand the relationship between variant sequence, structure and function in the ILPR we characterised both the C-rich (ILPRC, Table 1) and the G-rich sequences (ILPRG, Table 2) of the 11 main native variants. Each tandem repeat has two tracts of guanines or cytosines (5’-ACA<u>GGGG</u>TGT<u>GGGG</u>-3’/3’-TGT<u>CCCC</u>ACA<u>CCCC</u>-5’) so we designed our sequences to have two repeats, which would provide the minimum sequence necessary for i-motif or G-quadruplex formation. We maintained flanking sequences either side of the terminal C/G-tracts in line with the tandem repeat sequence. The general sequence for each variant was Flank-(C/G-tract)-Loop_1_-(C/G-tract)-Loop_2_-(C/G-tract)-Loop_3_-(C/G-tract)-Flank. Each C-rich and G-rich variant was characterised using circular dichroism (CD) to determine the overall topology, thermal difference spectroscopy (TDS) to characterise the type of structure in solution, and UV melting/annealing experiments to determine the thermal stability. For the C-rich sequences, the transitional pH was also determined by CD, to allow comparison of the pH stability of the sequence variants.

**Table 1:**
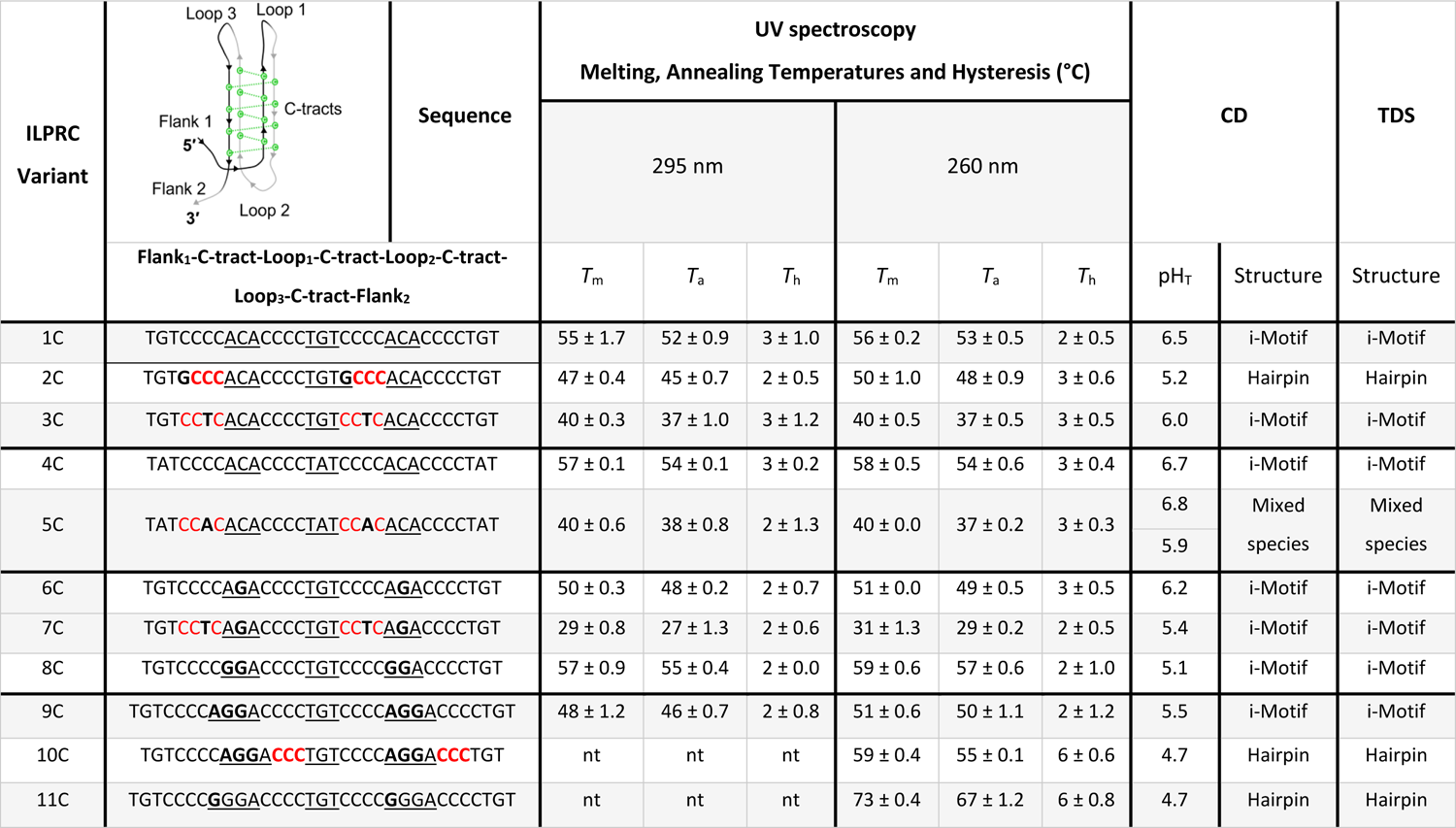
Sequences of the C-rich ILPR (ILPRC) variants - loops are underlined, mutations are in bold, and incomplete C-tracts are shown in red. Melting, annealing and hysteresis temperatures, transitional pH, and structure characterisation from the CD and thermal difference spectra (TDS). Data shown as mean ± SD (n= 3). Nt = no transition observed.

**Table 2:**
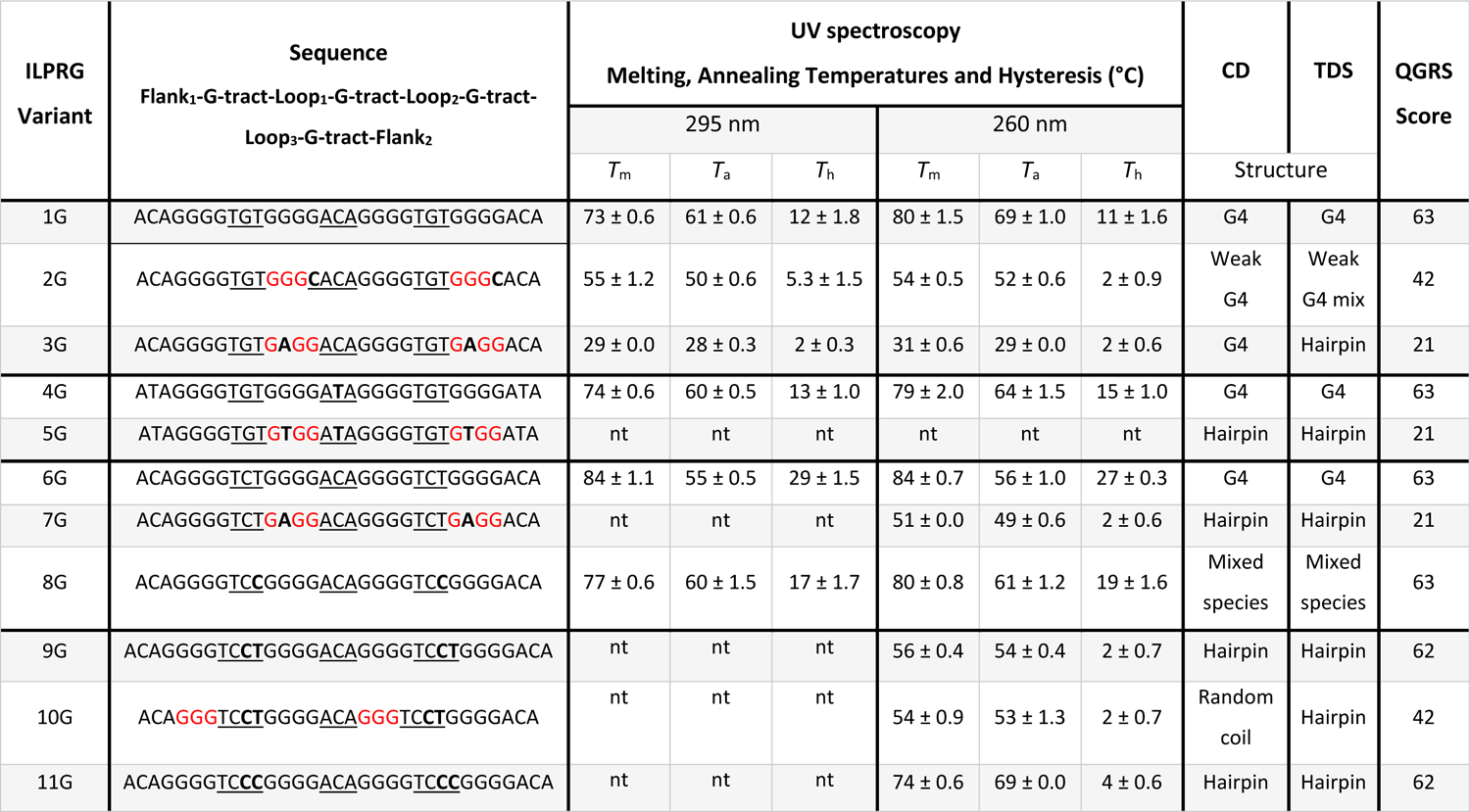
Sequences of the G-rich ILPR (ILPRG) variant sequences - loops are underlined, mutations are in bold and incomplete G-tracts are shown in red. Melting, annealing and hysteresis temperatures, transitional pH and structure characterisation from the CD and thermal difference spectra and QGRS score. Data shown as mean ± SD (n= 3). Nt = no transition observed.

The C-rich ILPR sequence variants were characterised in 10 mM sodium cacodylate buffer with 100 mM KCl. CD spectroscopy was performed at a range of pHs between 4 and 8 for determination of the transitional pH. TDS and UV melting and annealing experiments were performed at pH 5.5 to allow for assessment of the relative stability of all sequences, even those that may not be stable at physiologically relevant pH. We considered this pH would allow for more stable and less stable variants to be characterised fully and allow all sequences to be compared alongside each other. A summary of sequence, the melting temperature (*T*_m_), annealing temperature (*T*_a_), the thermal hysteresis (*T*_h_), transitional pH, and structural assignment by TDS are provided in Table 1. The corresponding example data is provided in the supporting information (Figures S1-5).

The predominant ILPR C-rich variant (1C) gave clear UV melting and annealing traces (Figure S1) and a *T*_m_ of 55 ± 1.7°C (Table 1). This melting temperature is higher than that measured by others for the same sequence, but the previous work was performed in phosphate buffer, which displays reduced buffering capacity at elevated temperatures.^20^ The TDS of sequence 1C showed positive peaks at 240 and 265 nm and a negative peak at 295 nm, consistent with i-motif structure (Figure 1A).^21^

**Figure 1:**
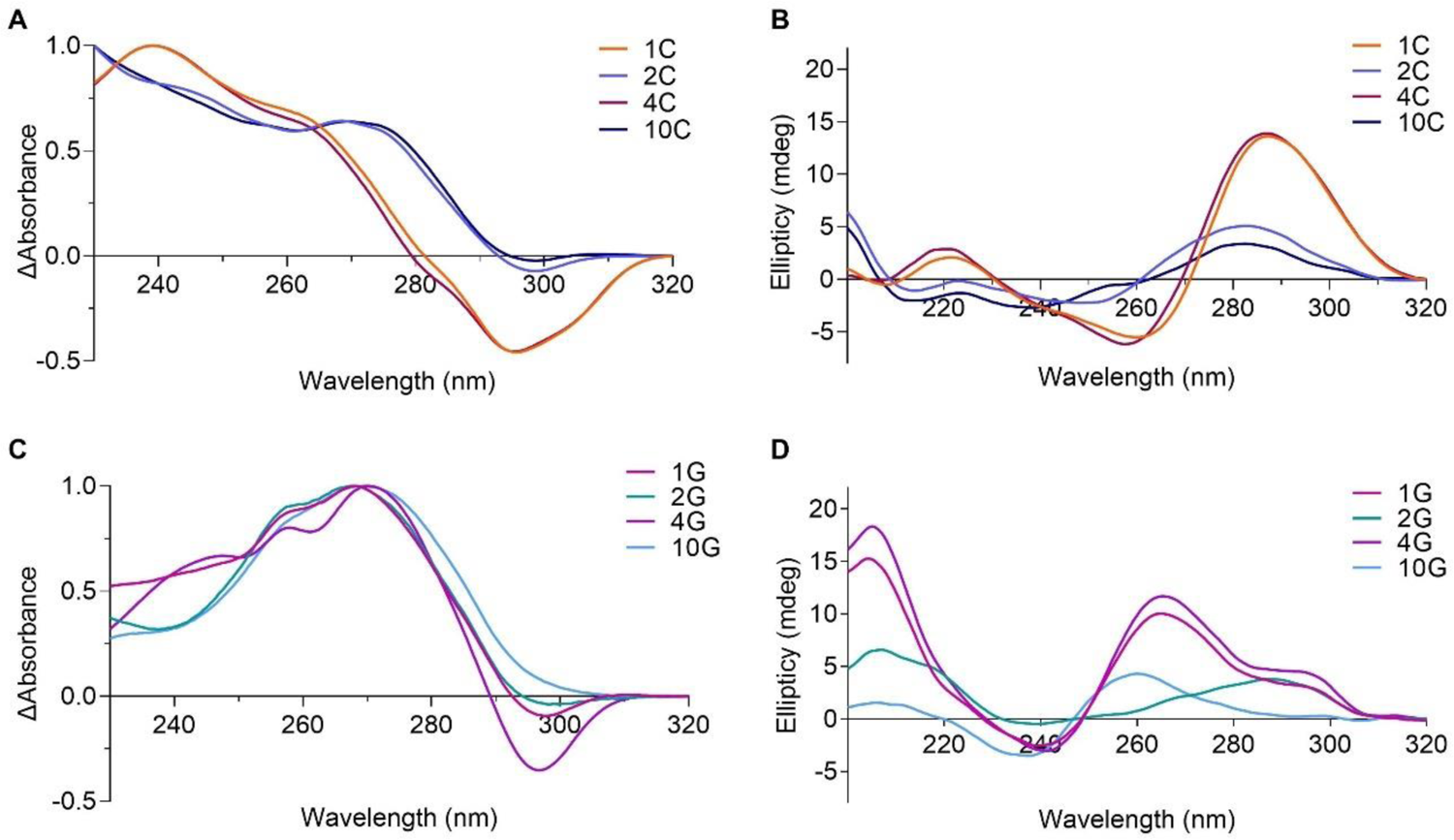
Example circular dichroism and thermal difference spectroscopy of ILPR Variants. A) TDS of 2.5 μM 1C, 2C, 4C and 10C in 10 mM sodium cacodylate, 100 mM KCl at pH 5.5. B) CD spectra of 10 μM ILPRC variants 1C, 2C, 4C and 10C in 10 mM sodium cacodylate, 100 mM KCl at pH 5.5. C) TDS of 2.5 μM ILPRG variants 1G, 2G, 4G and 10G in 10 mM sodium cacodylate, 20 mM KCl at pH 7.0. D) CD spectra of 10 μM ILPR variants 1G, 2G, 4G and 10G in 10 mM sodium cacodylate, 100 mM KCl at pH 7.0.

Similarly, CD spectroscopy of variant 1C showed i-motif formation at acidic pH, indicated by a positive peak at 288 nm and a negative peak at 260 nm (Figure 1B and S2).^22^ As the pH increases towards pH 7, the conformation unfolds, the positive peak shifts to 273 nm and the negative peak to 250 nm (Figure S2A). The transitional pH of 1C was determined to be 6.5 (Table 1 and Figure S2F), which was in-line with previous experiments for this sequence variant.^23^ One ILPRC variant (4C) demonstrated the same stability as sequence 1C, with a transitional pH of 6.7 (Table 1, Figure S2I) but the other variants were all significantly less pH stable, with transitional pHs as low as 4.7 (sequence variants 10C and 11C, Table 1 and Figure S4). Interestingly, minor differences in sequence made significant changes in the stability of the structures formed. For example, in ILPRC variant 2C, a single C to G mutation in each of the tandem repeats results in significantly lower melting (47 ± 0.4°C compared to 55 ± 1.7°C) and annealing temperatures (p<0.0001) and also a lower transitional pH of 5.1 (p<0.0001). It appears to be a hairpin-like structure at pH 5.5 in the TDS analysis (Figure 1A) and from the CD spectra at pH 5.5 (Figure 1B). I.e., this C to G mutation prevents i-motif formation.

Two main factors appear to decrease the stability of i-motif structure in these variants: mutation of loop nucleotides from cytosine to guanine (variants 6C-11C – highlighted in bold, in Table 1) or mutation/truncation within the C-tracts (variants 2C, 3C, 5C, 7C and 10C – highlighted in red, in Table 1). Some sequences are affected by both these factors (variants 2C, 7C and 10C, Table 1) and have some of the lowest transitional pHs overall (pH_T_s of 5.2, 5.4 and 4.7, respectively). The C to G mutation is clearly critical as it not only removes cytosines from the core stack of base pairs, but also introduces potential competing Watson/Crick complementary nucleotide which can shift the conformational equilibrium towards hairpin formation. This is further supported by the data acquired for sequences 10C and 11C which have more guanines in the loops. Both of these sequences did not give any transitions in the UV melting/annealing experiments at 295 nm (Figure S1D and S1L), but did so at 260 nm (Figure S1H and S1P), indicative of hairpin/duplex formation. TDS also indicated a spectrum inconsistent with i-motif and more consistent with that of duplex (Figure 1A and Figure S5), suggestive that these sequences in particular, form only hairpins under these experimental conditions. *In silico* structural calculations of these sequences using M-fold,^24^ also show clear potential for these sequences to form into hairpins (Figure S6). It is noteworthy that these sequences also demonstrate a larger hysteresis (6°C) compared to the other ILPRC variants (generally 2-3°C), indicative of slower kinetics in the formation of these hairpin structures.

Given the vast differences in structures and stability of the C-rich sequences, we wanted to also examine the complementary G-rich sequences, to see whether there was any complementarity in terms of the structures formed.

The G-rich ILPR sequence variants were also characterised by CD in 10 mM sodium cacodylate buffer, pH with 100 mM of KCl, NaCl or LiCl to give an indication of cation preference typically observed in G-quadruplex forming sequences. Sequences were characterised by UV melting/annealing in analogous buffer except with 20 mM KCl (100 mM concentrations of KCl resulted in *T*_m_ values >95°C). A summary of the characterisation of the sequences in KCl cation conditions: the melting temperature (*T*_m_), annealing temperature (*T*_a_), the hysteresis (*T*_h_), QGRS mapper score,^25^ and structural assignment by CD and TDS are provided in Table 2. The corresponding example data is provided in the supporting information (Figures S7-9).

The predominant ILPR G-rich variant (1G) gave clear UV melting and annealing traces (Figure S7A) and a *T*_m_ of 73 ± 0.6°C (Table 2). This melting temperature is similar to that measured previously (∼78°C) for the same sequence but in Tris buffer.^26^ The TDS of sequence 1G showed several positive peaks at 240, 255 and 270 nm and a negative peak at 295 nm, consistent with G-quadruplex structure^21^ (Figure 1C). Variant 1G gave CD spectra with a negative peak at 245 nm, and positive peaks at 263 nm and 295 nm (Figure 1D). This is in line with previously published CD spectroscopy data, showing a mixed population of parallel and antiparallel G-quadruplex formation in presence of KCl and a shift towards antiparallel G4 formation in the presence of weaker stabilizing cations NaCl and LiCl (Figure S8A).^14, 26, 27^

Of the other G-rich sequence variants, 4G (*T*_m_ = 74 ± 0.6°C) had similar thermal stability compared to 1G (73 ± 0.6°C), showing the mutation in G-tract Loop_2_ from a C to a T makes little difference in the stability (Figure S7B and S7F). ILPRG variant 8G was more stable (77 ± 0.6°C) than 1G, but does present as a potential mixture of species by CD and TDS (Figure S7K, S8G, and S9). The most stable of all the variants was sequence 6G (84 ± 1.1°C). 1G, 4G and 6G were all clearly characterized as G-quadruplexes by CD and TDS in KCl cationic conditions, similar to previously described studies on these sequences (Figure S8 and S9).^19, 27^ Some sequences formed significantly weaker secondary DNA structures and were thermally less stable than these four strong G-quadruplexes. For example, 2G, which is the reverse complement of 2C, has a significantly lower melting (*T*_m_ = 55 ± 1.2°C compared to the 1G variant with a *T*_m_ of 73 ± 0.6°C) and annealing temperatures (p<0.0001). This sequence (2G) also presents as a mixture of G-quadruplex and a hairpin-like structure in the TDS analysis (Figure 1C) and the CD spectra shows a broad weak positive peak at 300 nm and a negative peak at 245 nm (Figure 1D). This is in line with a formation of a weak antiparallel G-quadruplex, and potentially mixed with some sort of hairpin/duplex.^22^ The fact that there is a melting transition in the UV at 295 nm (Figure S7I) is potentially indicative of the former DNA structure, however, a negative peak at 295 nm in the TDS may present with Z-DNA and Hoogsteen DNA as well as G-quadruplex and i-motif structures.^22^ Interestingly, some of the sequences (5G, 7G, 9G, 10G, and 11G) do not have UV melt and anneal profiles at 295 nm, as expected with G-quadruplex structures. However, variants 7G, 9G, 10G, and 11G have clear melting and annealing transitions at 260 nm, consistent with these sequences forming hairpins or duplex-like structures (Figure S7).^28^ For example, the 10G variant, is similar to 2G in the TDS signature (Figure 1C), consistent with hairpin formation and clearly different to that of the G-quadruplexes formed by 1G, 4G, and 6G (Figure S8). The CD spectrum of 10G shows only a very weak positive signal at 260 nm and a negative signal at 240 nm (Figure 1D), which is consistent with unfolded G-rich sequence or a very weak G-quadruplex. Notably, the hairpin forming variants also lost cation sensitivity in the CD spectra and all have a narrow dip in signal at 215 nm (Figure S8). M-fold predictions of all tested variants show clear potential for these sequences to form into hairpins (Figure S10).

These results indicate that although particular ILPR variants are capable of forming i-motif and G-quadruplex structures, not all of them do. Comparing the biophysical data with the G-scores from QGRS mapper^25^ (Table 2) indicates that QGRS mapper accurately predicts the most stable G-quadruplex forming sequences 1G, 4G, 6G and 8G (all score 63), but there are two sequences that score nearly as high (9G and 11G, score 62) that do not form G-quadruplexes at all. Moreover, it is important to consider that sequences such as 2G and 10G do not form stable G-quadruplex structures, but score 42, i.e. the same score as the G-quadruplex forming sequence from the widely studied human telomere (TTAGGGTTAGGGTTAGGGTTAGGGTTA). Although the scores are only for G-rich sequences, this indicates that the nucleotide composition in the loops is critical for both C-rich and G-rich ILPR sequences, their stability, and the structures they can form. Taken together with the biophysical data for the C-rich and G-rich ILPR variants, it is clear that stable i-motifs are not exclusively formed in the complementary sequences of stable G-quadruplexes. For example, from the native variants 7/11 of the C-rich sequences form stable i-motif structures whereas only 3/11 of the G-rich sequences form a clear G-quadruplex structures. This indicates that it may be easier to mutate out a G-quadruplex based on the sequence, whereas an i-motif structure is more difficult to eliminate completely.

### i-Motif and G-quadruplex Structures Control Insulin Gene Transcription

From the biophysical data it was clear that not all native ILPR variants form i-motif or G-quadruplex structures. Importantly, the most common variants (1C and 4C) formed the most stable i-motif structures and the complementary strands (1G and 4G) also formed stable G-quadruplex structures. We hypothesised that the secondary DNA structures forming into i-motifs and G-quadruplexes in the ILPR are key elements to control transcription of insulin. To test this hypothesis, we compared four ILPR variants using a Luciferase-based reporter gene assay, where the entire human insulin promoter up to the start of the ILPR was cloned upstream of the gene for Firefly Luciferase. The resulting Firefly bioluminescence is proportional to the insulin promoter activation. Due to the difficulty in cloning long lengths of the ILPR and the fact that this region of DNA is intrinsically variable between people, we included enough repeat sequences to form one i-motif or G-quadruplex. We chose the most common variants (1C/1G and 4C/4G) which formed both stable i-motif and G-quadruplex structures and two of the variants that appeared to form hairpin structures in both C-rich and G-rich sequences (2C/2G and 10C/10G).

Functioning β-Cells normally secrete insulin in response to increased blood glucose levels as part of the homeostasis of blood glucose. There are many cell line models which can be used to assess levels of insulin expression *in vitro*. These cells retain normal regulation of glucose-induced insulin secretion, allowing use of glucose as a positive control.^29^ We selected the rat insulinoma-derived cell line INS-1 as model system due to the lack of an intrinsic ILPR or analogous sequence.^30, 31^ The INS-1 cells were co-transfected with either one of the ILPR vectors and a reference vector encoding *Renilla* Luciferase to allow normalisation of transfection efficiency variations between experiments. After transfection, the cells were starved overnight and were treated with either fresh low (2.8 mM) or high glucose (16.2 mM) medium to determine their respective responsiveness to glucose after four hours.

The four ILPR variants showed no significant difference in firefly luciferase expression in presence of low (2.8 mM) glucose levels (Figure 2). However, in the presence of high glucose (16.2 mM) medium, there was a significant increase in the expression of luciferase relative to the control for the 1C/G and 4C/G plasmid variants (where the underlying sequences were shown to form stable i-motif and G-quadruplex structures) and no significant change in expression for the 2C/G and 10C/G plasmid variants (characterised to form hairpin-like structures).

**Figure 2.**
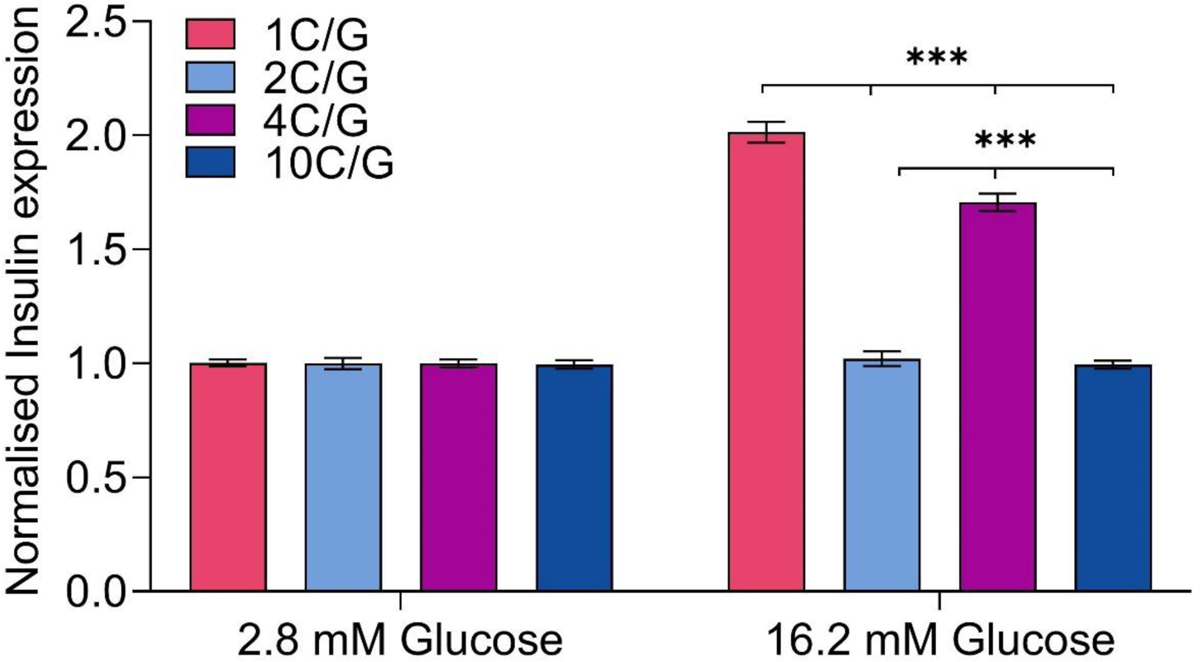
Dual Luciferase-reporter gene assay for glucose sensitivity in co-transfected INS-1 cells after four hours. Firefly signal is regulated by the human insulin promotor and is corrected to reference Renilla luciferase signal. Firefly to Renilla ratio is normalised to luminescence signals in low glucose levels, to represent insulin expression induced by glucose. Four different ILPR variants (1C/G, 2C/G, 4C/G or 10C/G) were measured. Mean ± SEM (n=12); p<0.001***, ns>0.1.

Specifically, the plasmid containing the 1C/G ILPR variant sequence showed a 2-fold increase in firefly luciferase expression levels (p<0.001) in the presence of high glucose levels compared to low glucose concentrations. It was expected for the most prevalent ILPR sequence (1C/G) to show glucose responsiveness and therefore we considered this as the positive control for this system. The plasmid containing the 4C/G ILPR variant sequence also responded to the higher glucose level with a 1.7-fold increase in gene expression compared to low glucose levels (p<0.001). Both example ILPR variants, with sequences capable of forming i-motif and G-quadruplexes responded to changes in glucose levels in a similar fashion, but the increase was significantly higher in the most prevalent ILPR variant (1C/G, p<0.001). Importantly, reporters encoding the two ILPR variants that did not form i-motif and G-quadruplex DNA structures (2C/G and 10C/G) showed no changes in the presence of 16.2 mM glucose (Figure 2). These data indicate the importance of the different sequence variants in the ILPR, showing the DNA structures they form control the responsiveness to glucose.

### Determination of the First Intramolecular i-Motif Crystal Structure

Given the apparent importance of the formation of i-motif and G-quadruplex structures in controlling relative insulin expression, we were interested in the potential interactions within the loops that made certain variants more stable than others. The most biologically relevant i-motif structure are intramolecular i.e. those formed from a single DNA strand, similar to what would form in the context of genomic DNA. However, structural information on intramolecular i-motifs is particularly scant. Although there are intramolecular NMR structures for i-motif (PDB IDs 1EL2, 1ELN),^32^ they are of modified fragments from the telomeres, and these modifications (necessary to enable structure determination by NMR) have been shown to alter the widths of the grooves in the structure.^33^ There are currently twelve intermolecular i-motif crystal structures formed from two or four separate strands but no intramolecular topologies. The apparent reason for the lack of intramolecular crystal structures is mainly due to the fact that i-motif loops are highly dynamic and difficult to resolve successfully using crystallographic methods. Intramolecular i-motif crystal structures would provide much opportunity for rational design of compounds to target these structures, and potential for drug development against these interesting biological targets, complementing recent drug discover projects targeting G-quadruplex.

With this in mind, we wanted to give the best chance for successful crystallisation, so we trialled the most stable C-rich ILPR variant that formed only i-motif from our biophysical studies: (4C) TATCCCC<u>ACA</u>CCCC<u>TAT</u>CCCC<u>ACA</u>CCCCTAT. This sequence is the second most prevalent ILPRC variant^13^ and lacks guanines within the sequence, so reduces the formation of intermolecular species through GC base pairing. To increase the chance of successful crystallisation we also designed different variants of this sequence with different flanking regions: TCCCCACACCCCTATCCCCACACCCCT (4Ca) and ATCCCCACACCCCTATCCCCACACCCC (4Cb) (Table S1). The crystallisations were performed at a pH below the pH_T_ (5.5), at which this sequence would be most stable.

Crystals of all three variants were obtained by hanging drop methods (Table S2) with the highest-quality diffraction data acquired with the original variant (4C). With the limited availability of i-motif structures, molecular replacement (MR) methods proved challenging. However, anomalous dispersion (AD) methods were successful for structure determination and model validation using both intrinsic and extrinsic scattering elements. Intrinsic phosphorous single – wavelength anomalous dispersion (P – SAD) where phosphorus is integral part of the DNA backbone provided validation of the native structural model (Figure S11) while the use of an extrinsic bromine, combined with multiple – wavelength anomalous dispersion (Br – MAD) methods provided anomalous scattering sufficient to generate high-quality maps for model building (Tables S3 and S4). In the 4C-Br sequence (Table S1), the Br substitution located within the less flexible CC core (Cytosine 4) provided a strong anomalous scattering contribution, while scattering for the second bromide in the middle loop (Adenine 16) was not observed due to the flexibility of this region.

The general use of intrinsic P anomalous scattering for structure determination of DNA/RNA motifs has proven challenging. The post analysis of our long wavelength data revealed a limited P anomalous scattering contribution, resulting in only a few P peaks of the anomalous difference map (*F*_anom(calc)_) overlapping the modelled positions or those observed with only weak diffuse peaks, this is despite using the lower energies closest to the peak (3.9995 Å, f’’2.3). The poor P-signal can be partly attributed to the static disorder, to the mobility of the phosphorous atoms and the low number of unique reflections compared to anomalous scatterers.^34^ We are currently exploring ways to optimise the P-signal for P-SAD applications.

### Structural Description of an Intramolecular i-Motif from the ILPR

The crystal structure formed from the ILPRC sequence 4C is comprised of two, independent, and inverted i-motifs in the asymmetric unit (Figure 3A and 3B). Each of these individual i-motifs is formed from four antiparallel strands held together by eight, intercalated hemi-protonated cytosine-cytosine base pairs, connected by three loops (Figure 3). The ACA loops connect strands at the minor grooves and the middle TAT loop at the major groove (Figure 3C, Table S5). The terminal CC-base pair is at the 3′-end, making each structure a 3′E topology (Figure 3B).^35^

**Figure 3.**
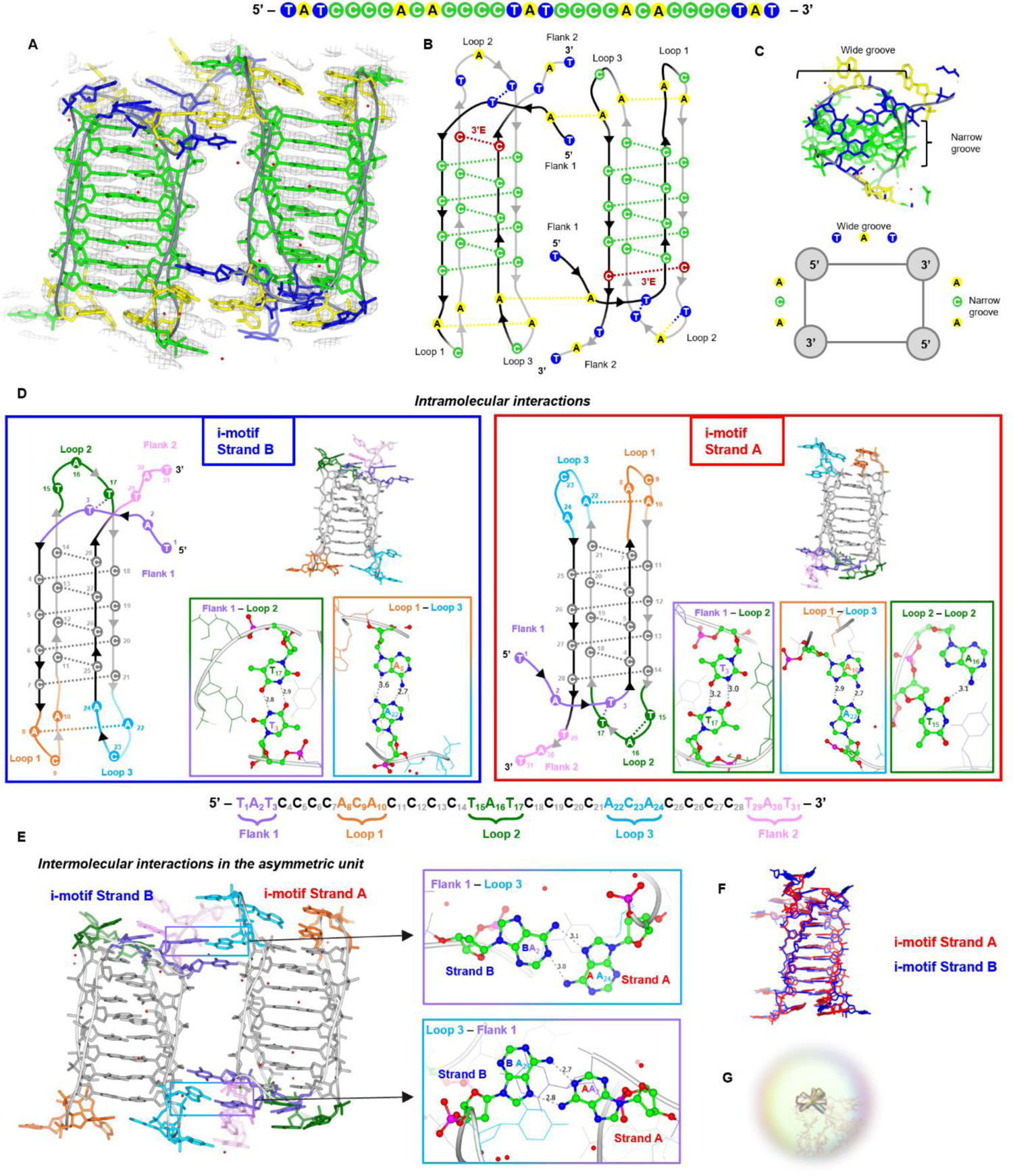
Crystal structure, structural features and interactions of the ILPR 4C intramolecular i-motif (PDB ID: 8AYG). (A) 4C structure coloured by nucleotide type (green: C, blue: T, yellow: A, grey: backbone, red: water molecules) and *2F_obs_ – F_calc_* electron density map contoured at 1.5 *σ* level (grey). (B) Schematic showing the two 4C intramolecular i-motifs as arranged in the asymmetric unit and the interactions they form. Both fold into a 3’E topology with the outer CC pair at the 3’–end (red). (C) Top view of one 4C i-motif and schematic showing the arrangement of the TAT and ACA loops at the wide and narrow grooves, respectively. (D) Intramolecular and (E) Intermolecular interactions formed by the two independent intramolecular 4C i-motif strands B and A present in the asymmetric unit. Each structure and schematic is coloured based on the flank or loop position in the sequence: flank 1 (purple), loop 1 (orange), loop 2 (dark green), loop 3 (light blue), flank 2 (pink), grey (C-core). The nucleotides involved in the hydrogen bonds shown in the boxes are coloured by atom type. All bond distances are in Å. (F) Structural comparison of the two i-motifs in the asymmetric unit by overlapping Strand A (red) and Strand B (blue). (G) Crystal of the 4C i-motif.

Apart from the CC base pairing, other interactions within each strand include mismatched base pairs like AA and TT which could contribute to the overall stability of the folded construct (Figure 3D, Table S6). In strand A, there is an AA base pair between loop 1 and loop 3, A_22_ (loop 3) interacts with A_10_ (loop 1) via two hydrogen bonds. The topology is further stabilized in loop 2, by the T_3_ from the flanking region, showing the importance of the flanking sequence in stabilizing interactions. Also, the flanking T_3_ interaction with T_17_ as a TT base pair stacks with the terminal CC-base pair (C_14_ and C_28_). T_15_ also forms a TA pair (T_15_ and A_16_) via one hydrogen bond, which then stacks on top of the TT base pair. While for strand B the TT base pair is sandwiched between the terminal CC base pair and an additional TAT triad consisting of T_15_ (loop 2), T_29_ (flank 2) and A_8_ (loop 1) from the symmetry of strand A (Figure S12, Table S7). Also, in strand B the AA base pair is formed between A_22_ (loop 3) and A_8_ rather than with A_10_ (loop 1) as in strand A.

Strand B is similar to strand A (Figure 3F) with an RMSD of 2.32 Å (when flanks are excluded - nucleotides 4 to 28). A difference is that the A_16_ is displaced with a symmetry-related adenine, but still stacks on top of the TT base pair. Also, A_8_ displaces A_10_ in the interaction with A_22_ which allows A_10_ to interact with a symmetry related thymine (Figure 3D, Figure S13). When only the core is included in the calculation, the RMSD is 1.04 Å showing the high similarity between the two cores. Differences in the torsion angles and sugar puckers of the two strands are shown in Table S8 and Figure S15 and are attributed to the phosphate backbone flexibility.

As there are two i-motifs in the asymmetric unit, this gives an excellent view as to how more than one i-motif may interact with each other like “beads on a string”. There are clear interactions between flank 1 of one strand (A_2_) and loop 3 (A_24_) of the other strand (Figure 3E). Also, there are various π-π stacking interactions between the outer nucleotides of the TAT flanks and an A or T in the loops which highlight the importance of the flanks in the crystal packing (Table S7). Intermolecular TA and CC pairs further contribute to the crystal packing (Figure S13, S14). Given the ILPR is a polymorphic region within the genome, comprised of tandem repeats, these intermolecular interactions are potentially important for consideration with how ligands and nuclear proteins may interact with these structures.

No specific hydration pattern was observed at the middle of the CC core as most of the cytosine hydrogen bond donors and acceptors are used in the formation of the CC pairs. Some waters at the major groove were seen hydrogen bonded with the H of N4 of the cytosine which is not involved in the CC base pairing and we observe a bridging with the phosphate O atoms. This is in agreement with some of the other tetramolecular or bimolecular i-motif crystal structures published e.g., 1CN0,^33^ 1BQJ,^33^ 8DHC, 8CXF,^36^ but no bifurcated hydrogen bond to O2 of a cytosine partner was seen. Based on the use of the *F*_anom(calc)_ maps (Figure S11) we can more confidently describe these peaks as water molecules and exclude sodium or chloride ions. Although limited by resolution, we do observe water molecules in the loops and these could represent potential sites for hydrogen bond interactions, something which will be useful in future ligand design or interactions with proteins. Given the potential binding pocket revealed by A_16_, which in strand A is base paired with T_15_ and in strand B this adenine was displaced with a symmetry-related adenine, this indicated to us that this site may also be interesting for potential targeting with ligands.

### Stabilising TT Base Pairs are Observed in Solution

Given the interesting additional base pairs within the crystal structure, we were interested whether these could be observed also in solution, so we performed NMR spectroscopy to examine the imino proton region. The imino proton region shows a set of peaks between 15.4 and 15.8 ppm, consistent with the presence of hemi-protonated cytosines (Figure S16).^15^ There are also additional imino proton signals at 10.9 and 11.5 ppm, consistent with the presence of TT base pairs.^37^ Importantly, there are no signals in the region between 12.5 and 14 ppm, where the imino proton signals from GC and AT base pairing would be expected.^37, 38^ An NMR annealing experiment (from 333 K to 277 K) revealed the formation of the CC base pairs first at 319 K, followed by the TT base pairs at 312 K (Figure S17). The evidence from the NMR indicates that the structure in solution is similar to that in the crystal structure, and the TT base pairs are weaker than the CC base pairs. A recent study looking at i-motifs using a DNA microarray containing 10,976 genomic i-motif forming sequences found that i-motifs with shorter loops (*n* = 1-4) had enhanced stability when the sequences had thymine residues directly flanking C-tracts.^39^ The presence of the TT base pairs in both the NMR experiments and the crystal structure provides structural evidence for the reason why this is the case.

### Enhanced Sampling Molecular Dynamics

To further explore the conformational landscape of i-motifs, we performed enhanced sampling molecular dynamics simulations. Of particular interest to us were the loops regions, which are the major contributor to the dynamics, also differentiate i-motifs from each other and from other nucleic acid structures; and the influence of the flanking nucleotides on the dynamics of the loops. Markov state models (MSMs) were built to study the kinetics of conformational transitions in the loop regions (for full details see the Supplementary Information, Figures S18-25 and Table S9). Upon creation of these models, both strands present a free energy landscape consisting of multiple metastable states that also explore the crystallographic conformations.

Given the interactions observed in the crystal structure originating from the flanking sequence, we looked at sequence 4C (TATCCCCACACCCCTATCCCCACACCCCTAT) and also an analogue with one base missing at the 3’-end, 4Cdel (TATCCCCACACCCCTATCCCCACACCCCTA).

Our analysis suggests that 4Cdel is far more dynamic with multiple interactions compared to 4C. Upon inspection of the structures extracted from the coarse-grained models, 4Cdel featured far more unstructured conformations in loops 1 and 3, while those in 4C seemed fairly ordered. Loop 2 in both sequences was well ordered. This would seem to suggest that slow motions in the i-motif structure are largely as a result of stabilising and compensating for the flexibility in not only loop 2 but also in the regions that flank it, namely the 5’ and 3’ ends. This can be visualised in the dynamics of 4C and 4Cdel.

The 4C structure is longer than 4Cdel by one base (T) at the 3’ end. This extra nucleotide leads to significantly more structural ordering via π-stacking interactions within the 3’ end itself, which then leads to the ordered conformations observed in loop 2. Comparing this with 4Cdel, the additional stacking is not possible and therefore the interactions with loop 2 produces a greater number of metastable states. Since time independent components (tICs) are ordered from slowest to fastest in terms of motions, those that provide the most stability will be ordered highest than those that are faster. But still a significant number may not be fully described by the number of dimensions which the features were reduced into. This is borne out by the unstructured conformations of loops 1 and 3 in these models as opposed to the fairly ordered ones of loop 2 and the terminal regions.

The simulation data supports the hypothesis that flanking sequences are important to the stability of i-motif structure, by providing the opportunities for additional interactions that reduce conformational dynamics. This is not only important for consideration of sequence designs for *in vitro* experiments involving i-motifs, but also may play an important role in how small molecules and proteins can interact with i-motif structures, and their consequential effects in biology.

## CONCLUSIONS

Here we show that different sequence variants of the ILPR form different DNA secondary structures and insulin expression is dependent on formation of i-motif and G-quadruplex structures. Importantly, not all native ILPR variants are capable of forming i-motifs and G-quadruplexes and minor changes in the sequence has been shown to give completely different secondary structures. The first crystal structure and dynamics of an intramolecular i-motif also reveals that sequences within the loop regions form additional stabilising interactions. These AA, TT and AT base pairs are critical to the formation of the stable i-motif structures that are required to control insulin expression and reveal pockets for rational based drug design. We also showed the importance of flanking sequence in the crystallisation of i-motif structures, through several intermolecular interactions in the crystal structure and supporting dynamics observed by enhanced sampling molecular dynamics. The outcomes of this work reveal the detail in formation of stable i-motif DNA structures, with potential for rational based drug design for compounds to target i-motifs from the ILPR.

## METHODS

### Oligonucleotides

All tested DNA sequences were synthesised and reverse phase HPLC purified by Eurogentec (Belgium) and prepared to a final concentration of 1 mM (biophysics) or 1.5 mM (crystallography) in ultra-pure water and confirmed using a Nanodrop. Each experimental section states the DNA concentration and buffer system in which they were prepared. Biophysical samples were annealed by heating for 5 mins at 95°C in a heating block and allowed to anneal by slow cooling to room temperature overnight. Crystallography samples were annealed using a thermocycler/PCR machine. The temperature was held at 60°C for 5 minutes, above the melting temperature for the sequences, followed by cooling to 51°C at a rate 1 °C/min and then to 4 °C with a rate 1°C/3 mins. A slower cooling rate of 1°C/3 mins was ideal to allow the sequences to fold and avoid gel formation which was observed when higher rates where used.

### Circular Dichroism Spectroscopy

The CD spectra of the selected ILPR sequences at different pH values, were recorded on a JASCO 1500 spectropolarimeter under a constant flow of nitrogen. The C-rich ILPR samples were diluted to 10 μM in 10 mM sodium cacodylate (NaCaco) and 100 mM KCl buffer at pH values ranging from 4 to 8. G-rich ILPR samples were prepared as 10 μM in 10 mM NaCaco and either in 100 mM KCl, 100 mM NaCl, or 100 mM LiCl buffer at pH 7.0. Four spectra scans were accumulated ranging from 200 nm to 320 nm for the buffer at each pH (blank) and DNA samples and measured at 20°C with a data pitch at 0.5 nm, scanning speed of 200 nm/min with 1 second response time, 1 nm bandwidth, and 200 mdeg sensitivity. Data was zero corrected at 320 nm and transitional pH (pH_T_) was determined for C-rich ILPR variants from the inflection point of the Boltzmann sigmoidal or bi-phase sigmoidal fit for the measured ellipticity at 288 nm and pH range.

### UV Melting/Annealing and Thermal Difference Spectroscopy

UV melting/annealing and TDS experiments were performed using the Jasco V-750 UV–Vis spectrometer. The C-rich ILPR samples were annealed at 2.5 μM in 10 mM NaCaco and 100 mM KCl buffer at pH 5.5 while 2.5 μM G-rich ILPR samples were annealed in 10 mM NaCaco and 20 mM KCl buffer at pH 7.0. For the UV melting/annealing experiments, the absorbance of the samples was recorded at every 1°C increase/decrease in three cycles at 295 nm and 260 nm. Initially, the samples were held at 4°C for 10 min followed by gradual increase to 95°C at a rate of 0.5°C/min (melting). When the temperature reached 95°C, it was held for 10 min before the process was reversed (annealing). The average melting (T_m_) and annealing temperature (T_a_) were identified by the first derivative method of for each measured cycle.

The thermal difference spectra (TDS) was obtained by measuring the absorbance spectrum from 230 nm to 320 nm after 10 mins at 4°C for the folded DNA structure and after 10 mins at 95°C for the unfolded structure. The TDS signature is determined by subtracting the absorbance spectra of the folded structure from the unfolded structure, zero corrected at 320 nm, and then normalised to the maximum absorbance.

### Crystallography Preparation of materials

The DNA sequences used in the crystallisations are provided in Table S1. Bromination in the 4C-Br sequence was on carbon-5 of the Cytosine_4_ and carbon-8 for the Adenine_16_. As we have previously shown, substitutions at these positions do not disrupt the folded i-motif topologies.^40^Oligonucleotides were annealed in the presence of 10 mM sodium cacodylate, 18 mM NaCl at pH 5.5.

### Crystallisations

Crystallisations were achieved using the hanging-drop vapor diffusion method. For the sequence 4C, the crystallisation solutions contained 49 mM sodium cacodylate buffer at pH 5.5, 45 mM NaCl, 23 % v/v 2-Methyl-2,4-pentanediol (MPD) and 2.9 mM spermine. For the drops, 1.2 μL of 0.3 mM 4C DNA solution and 1 μL crystallisation solution were combined and allowed to equilibrate. Pseudo-hexagonal crystals of the 4C sequence were formed in about two weeks at 10°C. It was still possible to grow good quality 4C crystals at 4°C but took longer. 4C-Br, 4Ca and 4Cb crystals were grown at 10 °C using the conditions summarized in Table S2.

### Data collection, phasing, and structure determination

Crystals were harvested using loops, placed in oil to remove mother liquor, and then cooled in liquid nitrogen. All diffraction data were collected at the Diamond Light Source synchrotron (DLS), UK.

P-SAD data for the 4C sequence were collected at the long wavelength I23 beamline in vacuum.^41^ The wavelength was tuned below the P-absorption edge (λ = 5.7788 Å, f’’4) to eliminate absorption. The maximum wavelength used during this experiment was 3.9995 Å, f’’2.3 to reduce absorption. Our data collections were undertaken at low-dose levels to prevent radiation damage and improve merging of multiple datasets. Good quality P-SAD data were collected but still resulted in a low P-anomalous signal as shown in Table S3. The signal from the intrinsic P atoms alone proved insufficient for phasing and structural determination, likely due to the low number of unique reflections compared to the number of anomalous scatterers.^34^ Based on data quality indicators, resolution range, completeness, *I / σ Ι* and R_meas_, the data collected at λ = 2.4797Å, f’’1.0 was selected consisting of six datasets merged to to increase the resolution and thus the number of unique reflections. Data were reduced using xia2.multiplex and subsequently re-scaled to 2.25 Å resolution with Aimless.^42, 43^ All data collection details are summarized in Table S3.

Data for the 4C–Br sequence were collected at the I03 beamline. A Br-MAD experiment was conducted at three wavelengths: 0.9196 Å (peak), 0.9203 Å (inflection point) and 0.9117 Å (remote). Data was reduced and scaled as described above. Data collection details are summarised in Table S4. The Shelx pipeline was used to determine the positions of the Br atoms and phase information.^44^ These phases were then used with the merged P-SAD dataset of 4C described above. Cycles of model building and refinement were then performed using COOT,^45^ and REFMAC5 (CCP4i package)^46^ with refinements within PHENIX (phenix.refine)^47, 48^ to generate a high quality model. This initial model was used to solve the structure of the native unmodified 4C sequence and refined to 2.25 Å resolution, the final R_free_/R_work_ values are 0.2948/0.2487. Refinement statistics are included in Table S3). Figures were generated with COOT and CCP4MG.^45, 49^

### Cell Culture

INS-1 rat insulinoma cells (AddexBio) were cultured in RPMI-1640 medium supplemented with 10% FBS and 50 µM 2-mercaptoethanol (BME), and 1% penicillin-streptomycin which were all obtained from Gibco. The medium was changed every four days and cells were expanded when reaching 80% confluency. Experiments were carried out in cells between passage 5-9.

### Transfection of Reporter Gene Plasmids

INS-1 cells were seeded at a density of 1 × 10^6^ cells per well in a 6-well plate in culture medium without antibiotics and transfected when 70-80% confluency is reached. The reporter plasmids were subcloned using the vector backbone of pGL410_INS421 which has a human insulin promotor regulating firefly luciferase expression (a gift from Kevin Ferreri Addgene plasmid #49057; http://n2t.net/addgene:49057; RRID:Addgene_49057). This plasmid had only 1.5 repeats of the predominant ILPR sequence (1C/G) which was sub cloned to replace the ILPR sequence by 2.5 repeats of 1C/G, 2C/G, 4C/G, or 10C/G. Each subcloned vector has unique flanks for selected restriction enzymes allowing confirmation of successful ILPR replacement via gel electrophoresis, also confirmed by sequencing. The reference vector pRL-TK Renilla plasmid was purchased from Promega UK Ltd and was co-transfected with each of the Firefly plasmids using Lipofectamine 2000 (Thermo Fisher). Each well of the 6-well plate was transfected with a total DNA amount of 3 µg with a 9:1 Firefly to Renilla ratio and a 1:1 DNA to Lipofectamine 2000 ratio, overnight. The following day, transfected cells and untreated INS-1 cells were seeded at a density of 1 × 10^5^ cells per well of a white 96 well plate in low glucose medium. The low medium was made up of RPMI 1640 medium (missing additives but L-glutamine and phenol red) supplemented with 2% FBS, 10 mM HEPES, 1 mM Sodium-pyruvate, 50 µM BME and 2.8 mM Glucose (all additives were obtained from Gibco). The overnight starved cells were treated with fresh low glucose medium or high glucose medium (16.2 mM). The four different ILPR variants were treated for 4h. The Dual Luciferase assay (Promega) was performed according to instruction manual and measured luminescence signals on SpectraMax iD3 (Molecular Devices). The resulting Firefly signal was corrected with the corresponding Renilla signal and final data was normalised to the Firefly/Renilla ratio of the low glucose signal to account for technical variation while preserve biological variation among 12 experimental repeats.^50^

Data analysis and presentation were performed using GraphPad Prism version 9.0. All sets of data passed all available normal distribution tests available in GraphPad Prism and presented as Mean ± SEM with indicated sample sizes (n). The statistical difference between treatments or variants was determined by one-way ANOVA and corrected with Holm-Sidak posthoc analysis.

### NMR

NMR data was recorded for the 4C ILPRC sequence. The DNA concentration was 0.66 mM in 9.1 mM sodium cacodylate buffer at pH 5.5, 91 mM KCl and 17% D_2_O. NMR data were acquired using a 700 MHz Bruker Avance III NMR spectrometer equipped with a TCI cryoprobe operating Topspin 3.6.2. 1H 1D spectra were acquired using a perfect echo watergate experiment,^51^ with a 1 s recycle delay, a 35 μs delay for binomial water suppression, a 780 ms acquisition time and a 30 ppm spectral width, and chemical shifts were referenced to the solvent. Data were processed with exponential window functions using nmrPipe.^52^

### Enhanced Sampling Molecular Dynamics

The initial ILPR i-motif crystal structure consists of two near-identical motifs packed into a dimer as a result of crystallisation. These two i-motifs were then separated into 4C (TATCCCC<u>ACA</u>CCCC<u>TAT</u>CCCC<u>ACA</u>CCCCTAT) and a modified shortened variant 4Cdel, which is missing the terminal T: (TATCCCC<u>ACA</u>CCCC<u>TAT</u>CCCC<u>ACA</u>CCCCTA). Adaptive Bandit simulations were run using the ACEMD molecular dynamics engine.^53^ Full details of the protocol are listed in the supplementary information. The simulation protocol was identical for both strands. The Markov State Models were built using the PyEMMA software.^54^

## DATA AVAILABILITY

Atomic coordinates and structure factors of the crystal structure have been deposited to the Protein Data bank under the identification code 8AYG.

## ACKNOWLEDGEMENTS

We thank Diabetes UK (DG and RVC, 18/0005820) and European Union’s Horizon 2020 research and innovation programme (ZD, project No 692068: BISON) for funding. We thank the DLS-CCP4 Data Collection and Structure Solution Workshop held in 2021 at the Diamond Light Source, UK and the teams at beamlines I23 and I03 who helped with the data collection and structure solution. NMR was supported by the Francis Crick Institute through provision of access to the MRC Biomedical NMR Centre. The Francis Crick Institute receives its core funding from Cancer Research UK (CC1078), the UK Medical Research Council (CC1078), and the Wellcome Trust (CC1078).

