## Supplementary Data for "Structural Insights into Regulation of Insulin Expression Involving i-Motif DNA Structures in the Insulin-Linked Polymorphic Region"

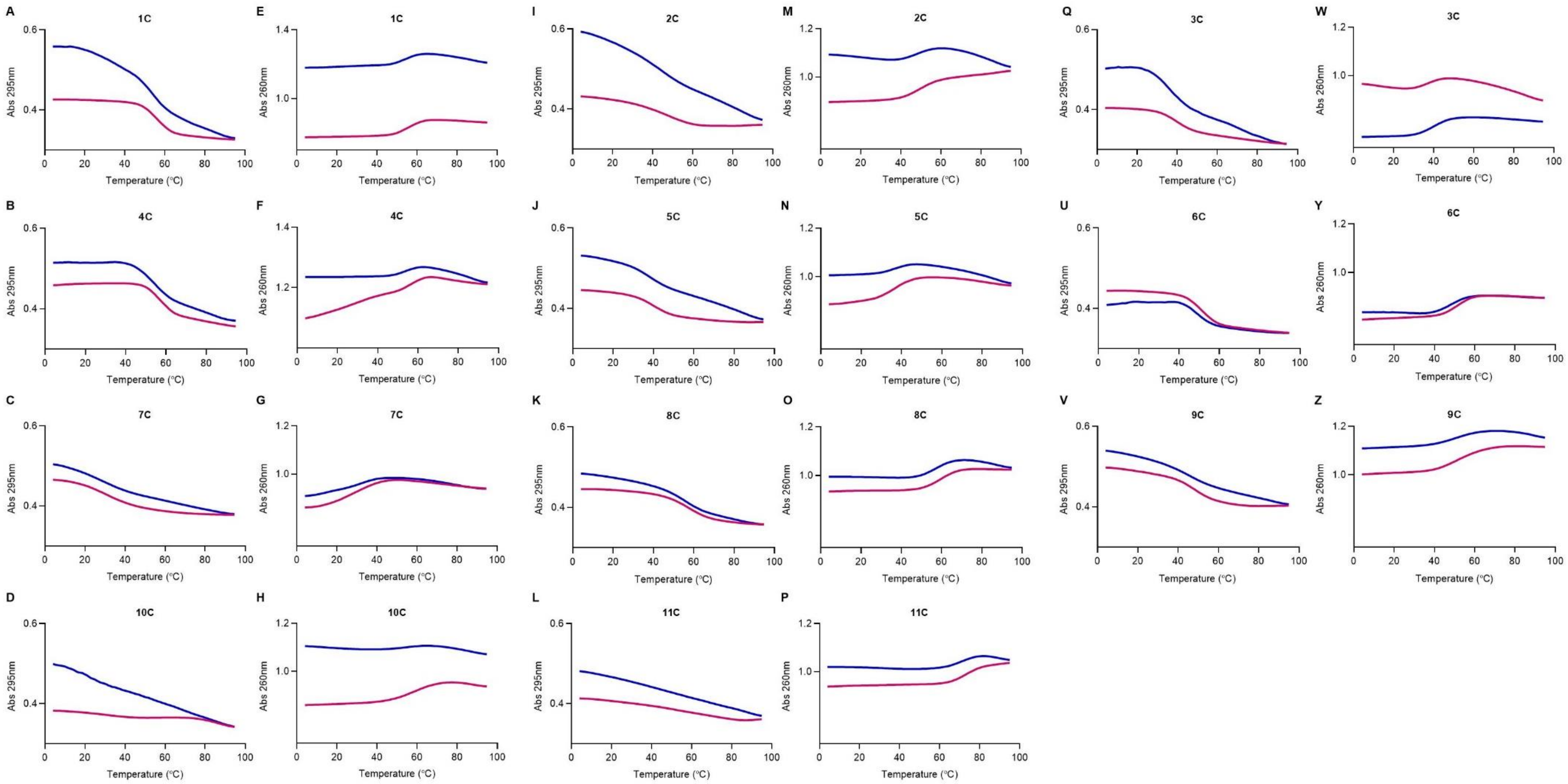


**Figure S1.** UV melting (purple) and annealing (blue) of C-rich ILPR sequences showing representative spectra at 295 nm (A-D, I-L, Q-V) and 260 nm (E-H, M-P, W-Z) of 2.5 µM DNA in 10 mM NaCaco 100 mM KCl at pH 5.5.

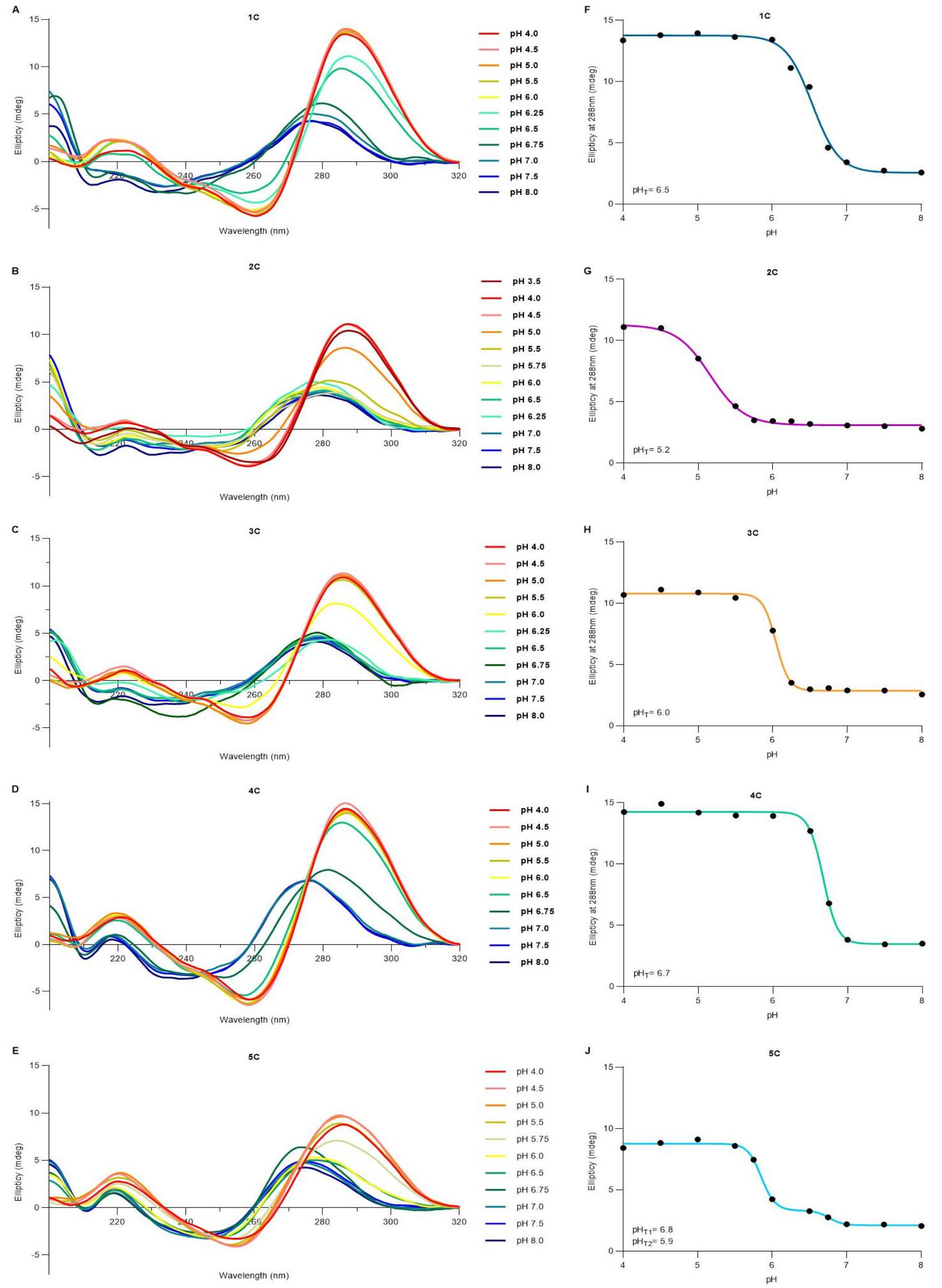


**Figure S2.** A-E) CD spectroscopy of C-rich ILPR sequences 1C, 2C, 3C, 4C, 5C. 10 µM DNA in 10 mM NaCaco 100 mM KCl and pH as indicated. F-I) Corresponding plot ellipticity at 288 nm at the different pHs to determine transitional pH ($pH_T$) from the inflection point of the Boltzmann sigmoidal or bi-phase fitted curve.

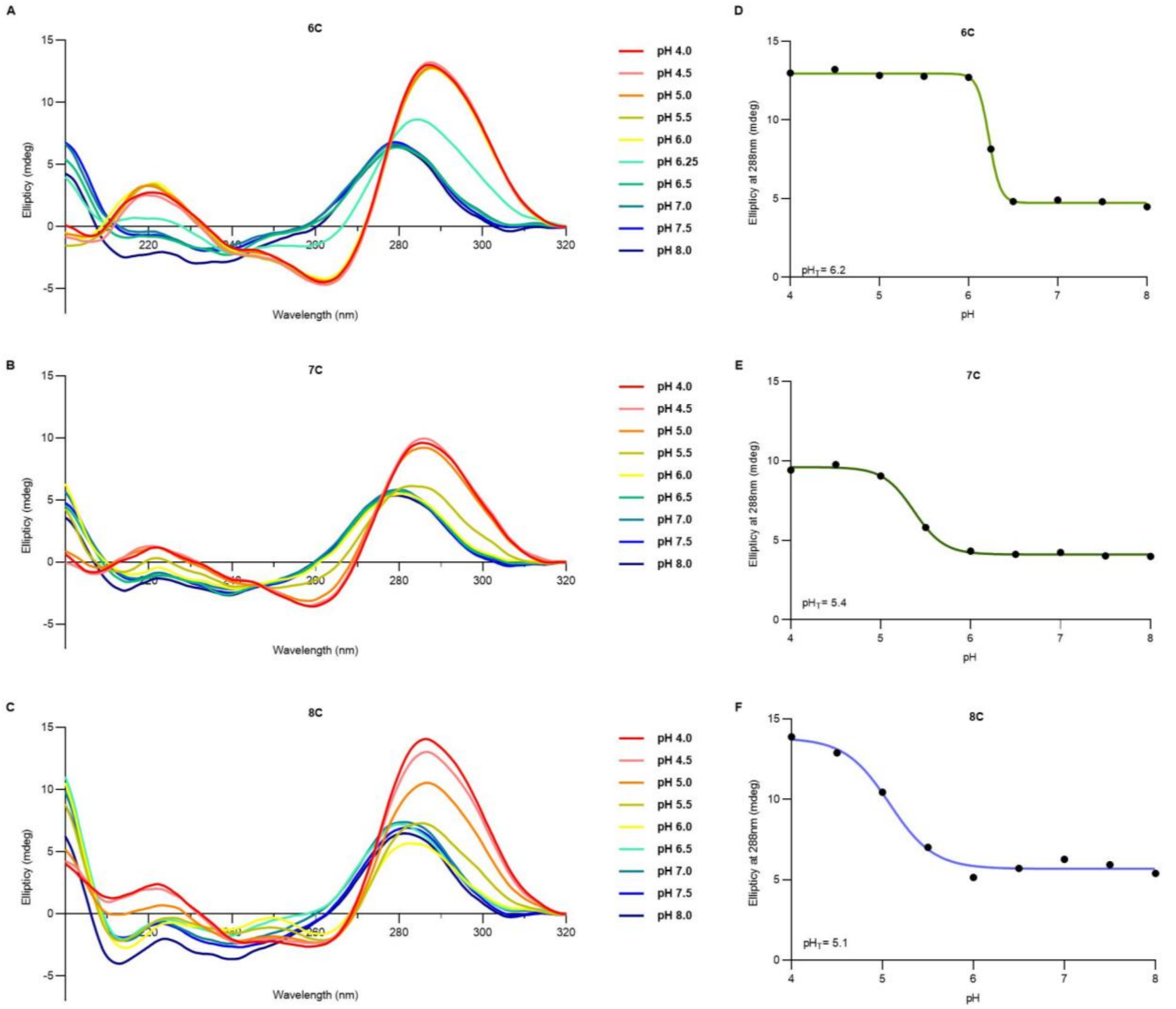


**Figure S3.** A-C) CD spectroscopy of C-rich ILPR sequences 6C, 7C, 8C. 10 µM DNA in 10 mM NaCaco 100 mM KCl and pH as indicated. D-F) Corresponding plot ellipticity at 288 nm at the different pHs to determine transitional pH ($pH_T$) from the inflection point of the Boltzmann sigmoidal or bi-phase fitted curve.

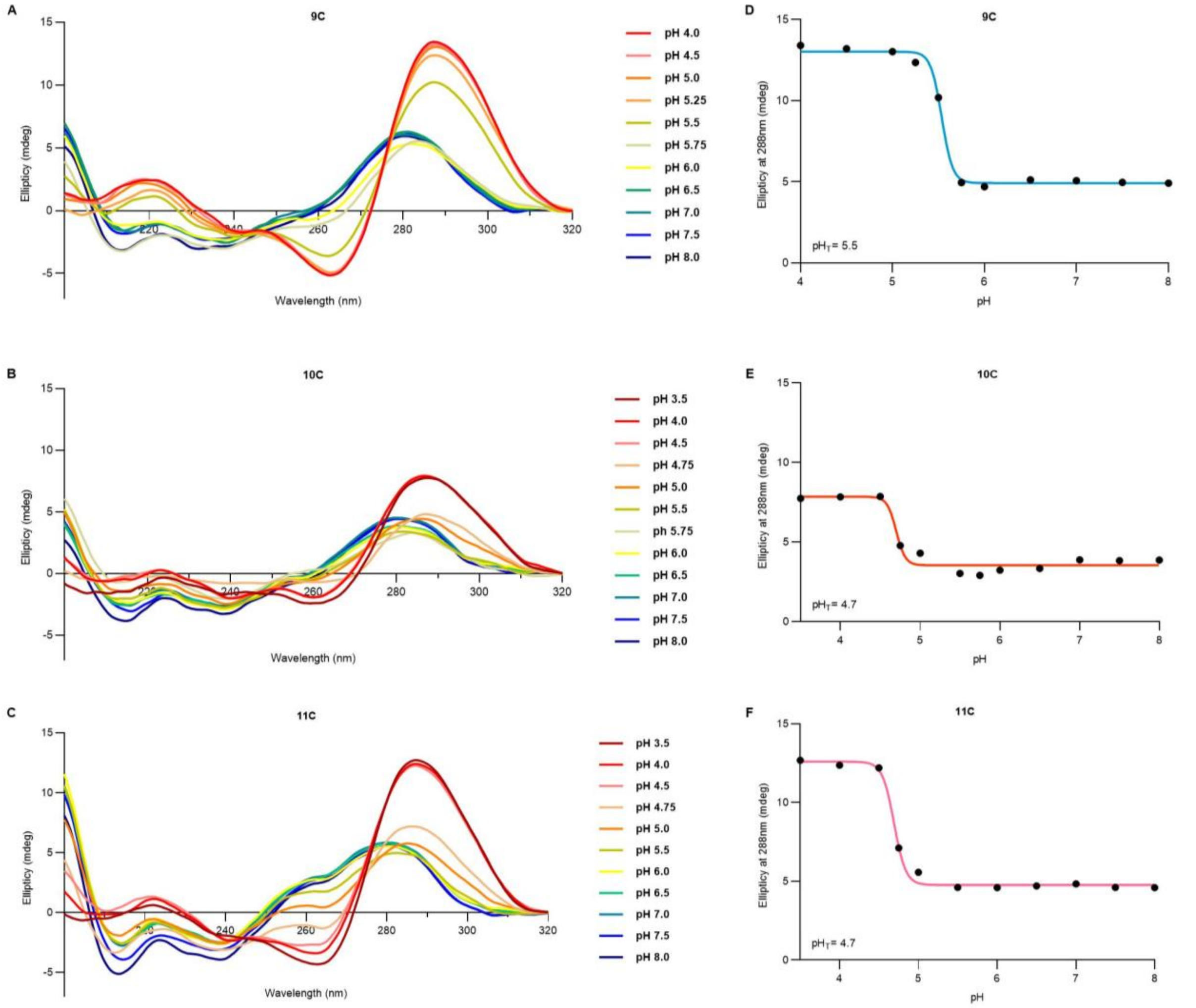


**Figure S4.** A-C) CD spectroscopy of C-rich ILPR sequences 9C, 10C, 11C. 10 µM DNA in 10 mM NaCaco 100 mM KCl and pH as indicated. D-F) Corresponding plot ellipticity at 288 nm at the different pHs to determine transitional pH ($pH_T$) from the inflection point of the Boltzmann sigmoidal or bi-phase fitted curve.

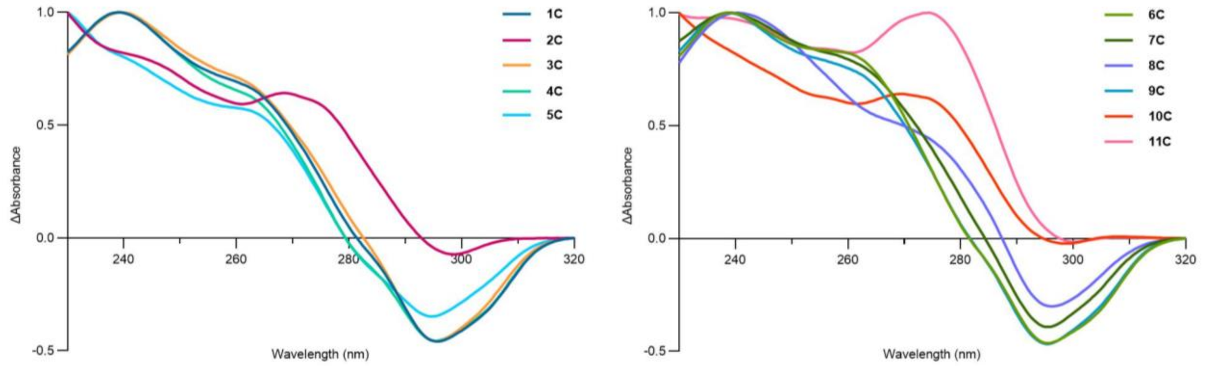


**Figure S5.** Thermal difference spectra of C-rich ILPR sequences with 2.5 µM DNA in 10 mM NaCaco 100 mM KCl at pH 5.5.

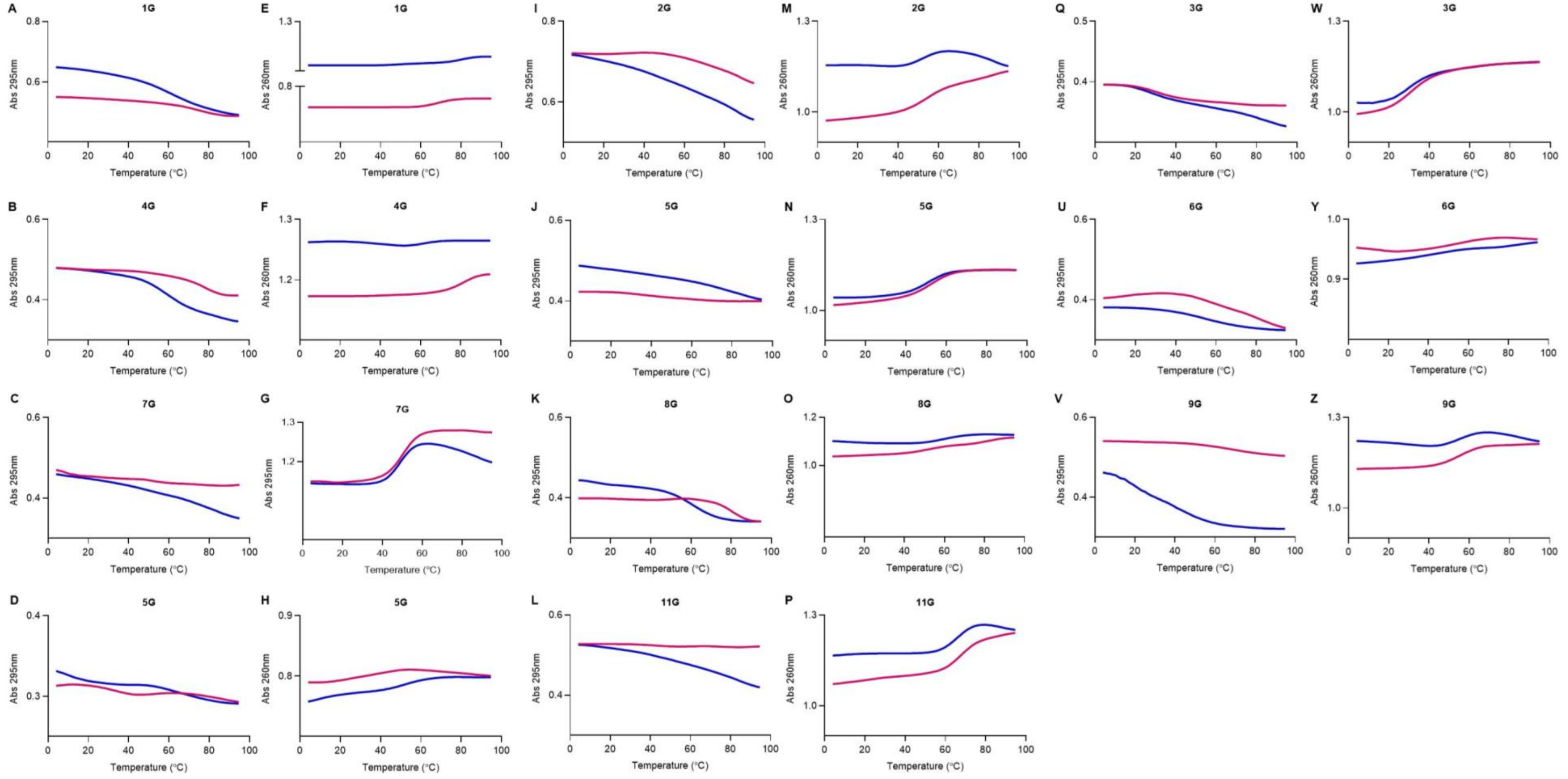


**Figure S6.** UV melting (purple) and annealing (blue) of C-rich ILPR sequences showing representative spectra at 295 nm (A-D, I-L, Q-V) and 260 nm (E-H, M-P, W-Z) of 2.5 µM DNA in 10 mM NaCaco 100 mM KCl at pH 5.5.

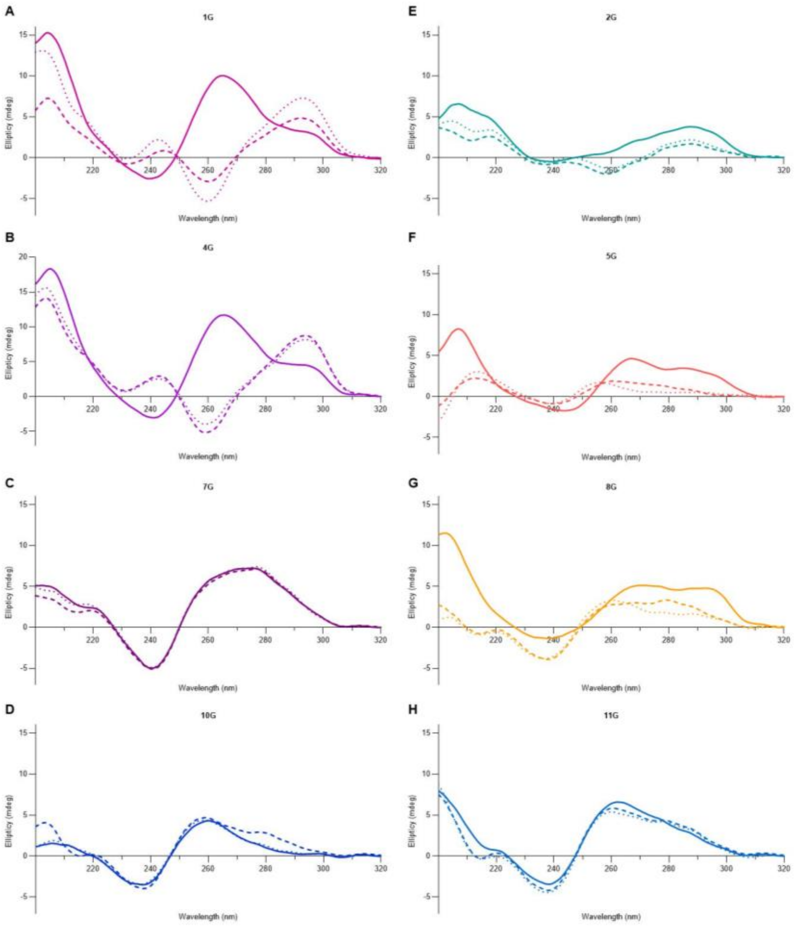


**Figure S7.** Biophysical characterisation of G-rich ILPR sequences via CD spectroscopy using 10 µM DNA in 10 mM NaCaco 100 mM KCl at pH 7.0 (solid line), 10 mM NaCaco 100 mM LiCl at pH 7.0 (dashed lined), or 10 mM NaCaco 100 mM NaCl at pH 7.0 (dotted line).

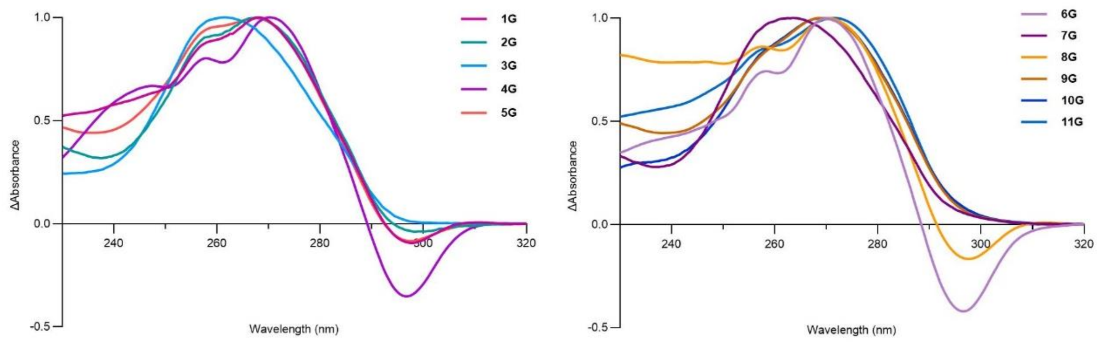


**Figure S8.** Thermal difference spectra of G-rich ILPR sequences with 2.5 µM DNA in 10 mM NaCaco 20 mM KCl at pH 7.0.

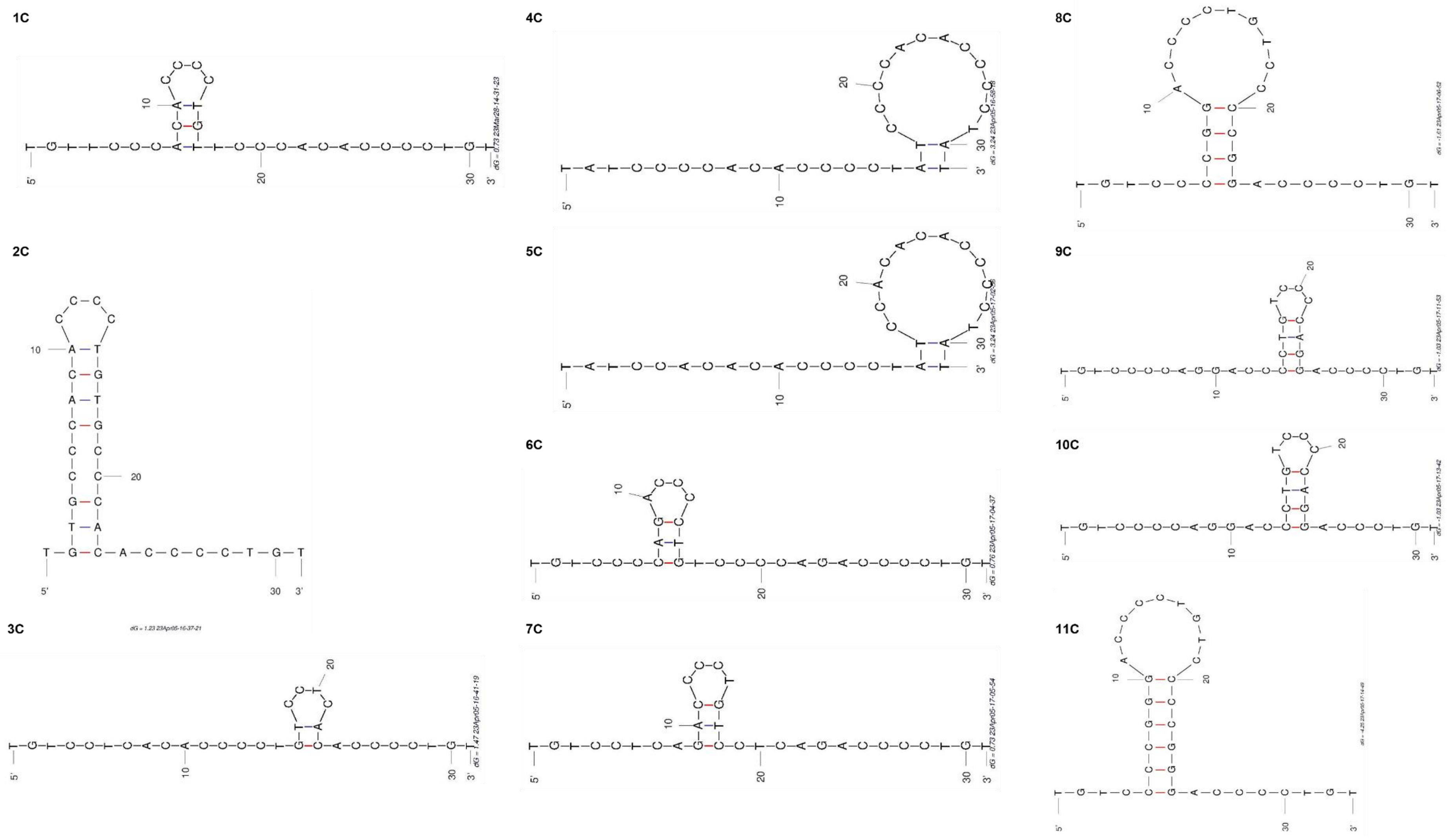


**Figure S9:** M-fold predictions of secondary DNA structures with highest predicted thermal stability from C-rich ILPR sequences.[1]

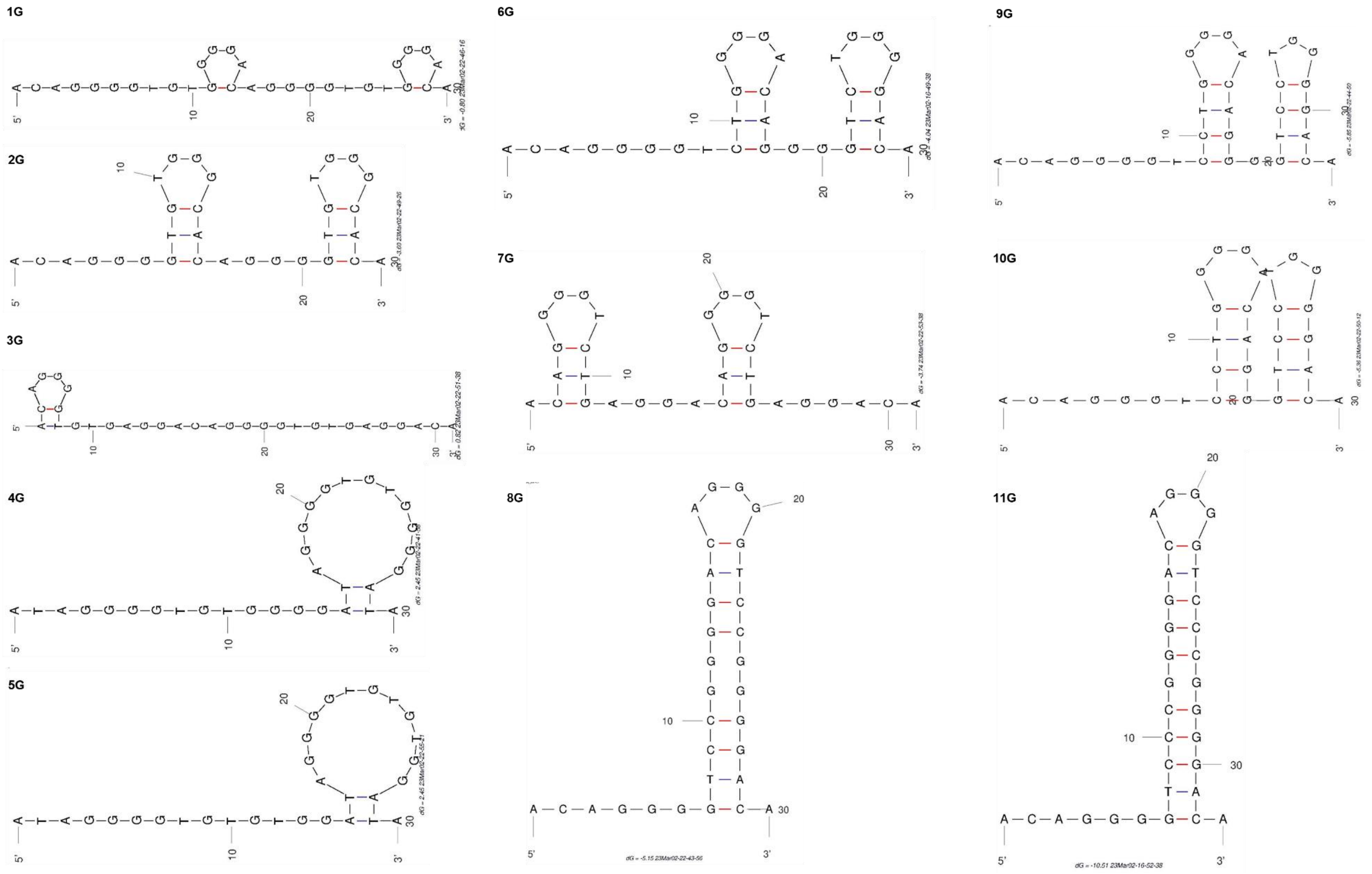


**Figure S10:** M-fold predictions of secondary DNA structures with highest thermal stability of G-rich ILPR sequences.

**Table S1.** The sequences of the 4C analogues with different flanking sequences and brominated sequence used for crystallisation screenings.

| 4C analogues | Sequence (5' → 3') |
|---|---|
| **4C** | TATCCCCACACCCCTATCCCCACACCCCTAT |
| **4Ca** | TCCCCACACCCCTATCCCCACACCCCT |
| **4Cb** | ATCCCCACACCCCTATCCCCACACCCC |
| **4C-Br** | TAT$^{Br}$CCCCACACCCCT$^{Br}$ATCCCCACACCCCTAT |

**Table S2.** Crystallisation conditions for obtaining crystals of each sequence.

| Sequence name | [DNA solution] (mM) | Crystallisation solution components | Drop: (µL DNA solution + µL crystallisation solution) | Crystallisation solution equilibrated against |
|---|---|---|---|---|
| 4C | 0.3 | 49 mM sodium cacodylate pH 5.5<br>45 mM NaCl<br>23 % v/v MPD<br>2.9 mM spermine | 1.2 + 1 | 49 mM sodium cacodylate pH 5.5<br>45 mM NaCl<br>23 % v/v MPD<br>2.9 mM spermine |
| 4C - Br | 0.3 | 49 mM sodium cacodylate pH 5.5<br>45 mM NaCl<br>23 % v/v MPD<br>2.9 mM spermine | 1+1 | 49 mM sodium cacodylate pH 5.5<br>45 mM NaCl<br>20 % v/v MPD<br>2.9 mM spermine |
| 4Ca | 0.75 | 49 mM sodium cacodylate pH 5.5<br>45 mM NaCl<br>2.2 M ammonium sulfate<br>2.9 mM spermine<br>10% v/v glycerol<br><br>Or<br><br>49 mM sodium cacodylate pH 5.5<br>45 mM NaCl<br>23 % v/v MPD<br>2.9 mM spermine | 1+1 | 49 mM sodium cacodylate pH 5.5<br>45 mM NaCl<br>2.2 M ammonium sulfate<br>2.9 mM spermine<br>10% v/v glycerol<br><br>Or<br><br>49 mM sodium cacodylate pH 5.5<br>45 mM NaCl<br>15 % v/v MPD<br>2.9 mM spermine |
| 4Cb | 0.75 | 49 mM sodium cacodylate pH 5.5<br>45 mM NaCl<br>23 % v/v MPD<br>2.9 mM spermine | 1+1 | 49 mM sodium cacodylate pH 5.5<br>45 mM NaCl<br>17-21 % v/v MPD<br>2.9 mM spermine |

**Table S3.** Data collection and refinement statistics for 4C.

Values in parentheses are for the highest-resolution shell.

| **Sequence name** | **4C** |
|---|---|
| **Sequence** | TATCCCCACACCCCTATCCCCACACCCCTAT |
| **Data collection** | |
| **Beamline** | Diamond Light Source I23 |
| **Wavelength (Å)** | 2.4797 |
| **Space group** | $P2_12_12_1$ |
| **Unit cell dimensions**<br>***a, b, c (Å)***<br>***α, β, γ (°)*** | <br>47.41 51.51 69.85<br>90.00 90.00 90.00 |
| ***I / σ I*** | 30.40 (1.7) |
| **$R_{meas}$ (within I+/I-)**<br>**$R_{meas}$ (all I+ & I-)** | 0.165 (2.009)<br>0.169 (2.014) |
| **$R_{merge}$ (within I+/I-)**<br>**$R_{merge}$ (all I+ & I-)** | 0.162 (1.920)<br>0.167 (1.966) |
| **Completeness (%)** | 95.2 (88.0) |
| **Multiplicity** | 44.5 (22.7) |
| **Resolution range (Å)** | 47.42 – 2.25 (2.32 – 2.25) |
| **Total number of reflections** | 362297 (14886) |
| **No. of unique reflections** | 8133 (657) |
| **$CC_{1/2}$** | 0.991 (0.748) |
| **Anomalous completeness %** | 94.6 (86.5) |
| **Anomalous multiplicity** | 23.3 (11.9) |
| **DelAnom correlation between half-sets** (inner shell) | 0.537 |
| **Refinement** | |
| **Completeness (working + test) (%)** | 93.14 (89.08) |
| **Resolution range (Å)** | 41.46-2.25 (2.58-2.25) |
| **No. of unique reflections refined against** | 7957 (2331) |
| **No. of molecules in the ASU** | 2 |
| **$R_{free}/R_{work}$** | 0.2948/0.2487 (0.4622/0.3748) |
| **No. of atoms**<br>**DNA (in the asymmetric unit)**<br>**DNA (per chain)**<br>**Water (in the asymmetric unit)** | <br>1212<br>606<br>22 |
| **Mean B value (overall Å$^2$)** | 80.06 |
| **RMSD from ideal values**<br>**Bond lengths (Å)**<br>**Bond angles (°)** | <br>0.013<br>1.113 |

**Table S4.** Data collection and processing statistics for 4C-Br.

Values in parentheses are for the highest-resolution shell.

| **Sequence name** | **4C – Br** | | |
|---|---|---|---|
| **Sequence** | 5' – TAT**$^{Br}$C**CCCACACCCCT**$^{Br}$A**TCCCCACACCCCTAT – 3' | | |
| **Beamline** | Diamond Light Source I03 | | |
| **Wavelength (Å)** | 0.9196 | 0.9203 | 0.9117 |
| **Space group** | $P2_12_12_1$ | $P2_12_12_1$ | $P2_12_12_1$ |
| **Unit cell dimensions**<br>***a, b, c* (Å)**<br>***α, β, γ* (°)** | 47.18 51.57 69.94<br>90.00 90.00 90.00 | 47.19 51.57 69.89<br>90.00 90.00 90.00 | 47.19 51.58 69.87<br>90.00 90.00 90.00 |
| ***I / σ I*** | 18.7 (2.6) | 21.9 (3.6) | 22.3 (3.9) |
| **$R_{meas}$ (within I+/I-)**<br>**$R_{meas}$ (all I+ & I-)** | 0.065 (1.085)<br>0.067 (1.083) | 0.058 (0.764)<br>0.058 (0.763) | 0.058 (0.697)<br>0.062 (0.697) |
| **$R_{merge}$ (within I+/I-)**<br>**$R_{merge}$ (all I+ & I-)** | 0.060 (1.008)<br>0.064 (1.043) | 0.054 (0.709)<br>0.055 (0.734) | 0.054 (0.646)<br>0.060 (0.670) |
| **Completeness (%)** | 98.9 (98.0) | 98.9 (98.2) | 98.9 (98.3) |
| **Multiplicity** | 13.2 (13.7) | 13.2 (13.7) | 13.2 (13.7) |
| **Resolution range (Å)** | 51.57 – 2.38<br>(2.47 – 2.38) | 51.57 – 2.38<br>(2.47 – 2.38) | 51.58 – 2.38<br>(2.47 – 2.38) |
| **Total number of reflections** | 94750 (10092) | 94731 (10133) | 94770 (10158) |
| **No. of unique reflections** | 7152 (734) | 7152 (738) | 7153 (740) |
| **$CC_{1/2}$** | 1.000 (0.897) | 1.000 (0.938) | 1.000 (0.857) |
| **Anomalous completeness %** | 99.3 (98.5) | 99.3 (98.6) | 99.3 (98.6) |
| **Anomalous multiplicity** | 7.2 (7.3) | 7.2 (7.3) | 7.3 (7.3) |
| **DelAnom correlation between half-sets** (inner shell) | 0.808 | 0.351 | 0.920 |

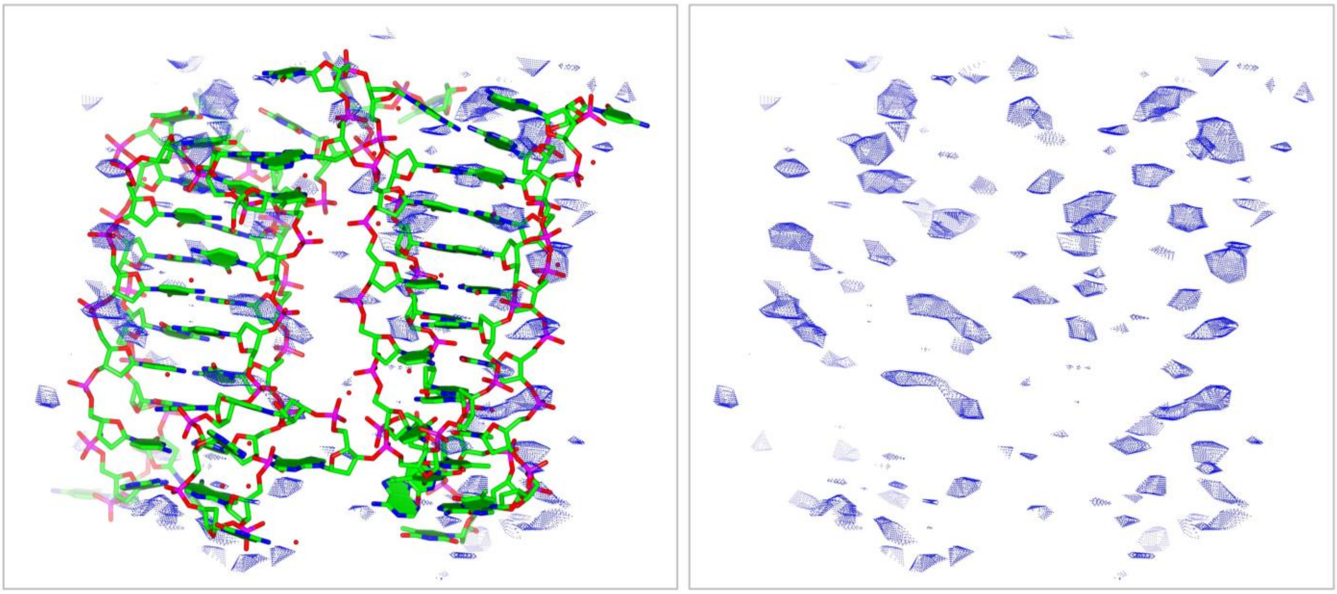

**Figure S11.** Anomalous difference map ($F_{anom(calc)}$, blue, 2$\sigma$) showing P atoms positions, 3.1 keV data (λ = 3.9995 Å).

**Table S5.** Narrow and wide groove dimensions based on P – P distances of adjacent strands. Average dimensions for each groove are shown in the schematic bellow with P – P distances of nucleotides from the cytosine rich core, loop or flank regions included in the calculation. The average dimension of the P-P distances at the core only was also calculated and shown below.

| Groove | Nucleotides | P – P distance (Å) | | Region |
|---|---|---|---|---|
| | | Strand A | Strand B | |
| Narrow groove 1 | $C_4$ – $T_{15}$ | 9.23 | 8.25 | Core – Loop 2 |
| | $C_5$ – $C_{14}$ | 6.32 | 6.58 | Core |
| | $C_6$ – $C_{13}$ | 7.20 | 7.06 | Core |
| | $C_7$ – $C_{12}$ | 6.97 | 6.73 | Core |
| | $A_8$ – $C_{11}$ | 6.87 | 7.65 | Loop 1 – Core |
| | $C_9$ – $A_{10}$ | 5.88 | 6.17 | Loop 1 |
| Narrow groove 2 | $C_{18}$ – $T_{29}$ | 7.64 | 6.52 | Core – Flank 2 |
| | $C_{19}$ – $C_{28}$ | 6.60 | 8.39 | Core |
| | $C_{20}$– $C_{27}$ | 6.96 | 6.76 | Core |
| | $C_{21}$ – $C_{26}$ | 5.80 | 5.98 | Core |
| | $A_{22}$ – $C_{25}$ | 9.36 | 7.98 | Loop 3 – Core |
| | $C_{23}$ – $A_{24}$ | 5.45 | 6.96 | Loop 3 |
| Wide groove 1 | $A_{10}$ – $A_{22}$ | 14.37 | 21.10 | Loop 1 – Loop 3 |
| | $C_{11}$ – $C_{21}$ | 12.20 | 13.41 | Core |
| | $C_{12}$ – $C_{20}$ | 12.35 | 14.03 | Core |
| | $C_{13}$ – $C_{19}$ | 12.53 | 13.50 | Core |
| | $C_{14}$ – $C_{18}$ | 12.50 | 13.75 | Core |
| | $T_{15}$ – $T_{17}$ | 11.31 | 12.13 | Loop 2 – Loop 2 |
| Wide groove 2 | $C_4$ – $T_{29}$ | 17.96 | 18.05 | Core – Flank 2 |
| | $C_5$ – $C_{28}$ | 17.06 | 16.56 | Core |
| | $C_6$ – $C_{27}$ | 16.47 | 16.66 | Core |
| | $C_7$– $C_{26}$ | 16.55 | 16.54 | Core |
| | [illegible] | [illegible] | [illegible] | Loop 1 – Core |

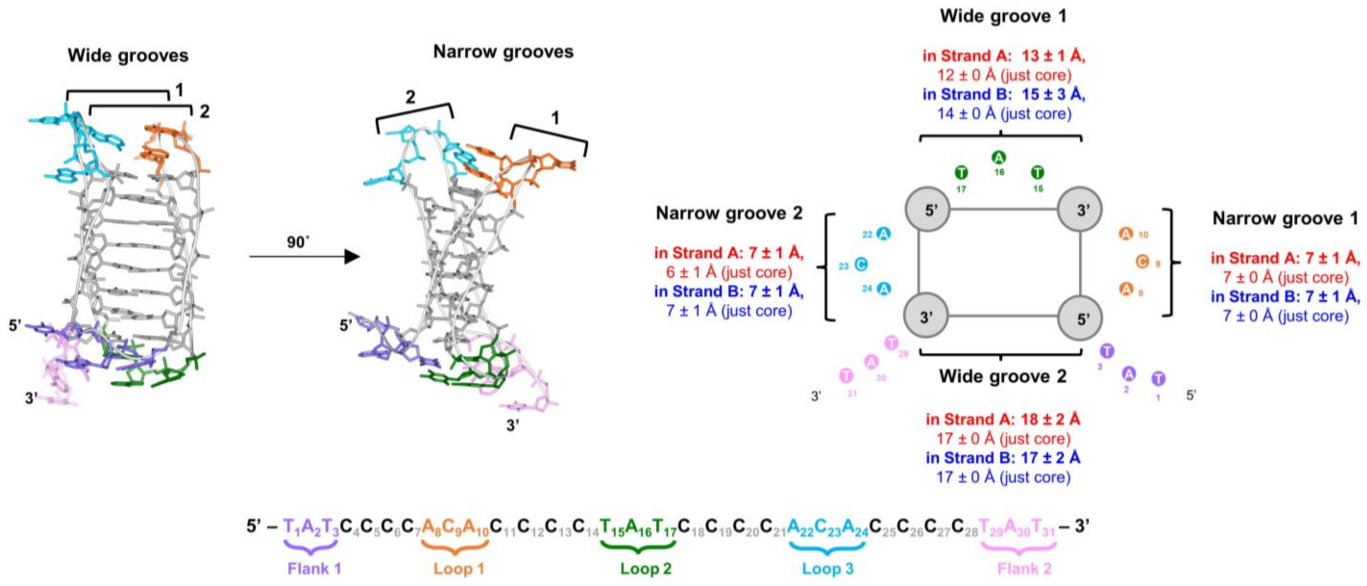

**Table S6.** Intramolecular interactions (within each strand) of the 4C i-motif based on the crystal structure.

5' – $T_1A_2T_3C_4C_5C_6C_7A_8C_9A_{10}C_{11}C_{12}C_{13}C_{14}T_{15}A_{16}T_{17}C_{18}C_{19}C_{20}C_{21}A_{22}C_{23}A_{24}C_{25}C_{26}C_{27}C_{28}T_{29}A_{30}T_{31}$ – 3'

Flank 1 ($T_1A_2T_3$) Loop 1 ($A_8C_9A_{10}$) Loop 2 ($T_{15}A_{16}T_{17}$) Loop 3 ($A_{22}C_{23}A_{24}$) Flank 2 ($T_{29}A_{30}T_{31}$)

| Mismatched TT and AA base pairs | | | |
|---|---|---|---|
| Strand | Nucleotides and atoms involved | Distance (Å) | Type of interaction |
| A | $T_{17}$ ($N_3$) Loop 2 – $T_3$ ($O_4$) Flank 1<br>$T_{17}$ ($O_4$) Loop 2 – $T_3$ ($N_3$) Flank 1 | 3.2<br>3.0 | Hydrogen bonds |
| | $A_{10}$ ($N_6$) Loop 1 – $A_{22}$ ($N_1$) Loop 3<br>$A_{10}$ ($N_7$) Loop 1 – $A_{22}$ ($N_6$) Loop 3 | 2.7<br>2.9 | Hydrogen bonds |
| B | $T_{17}$ ($N_3$) Loop 2 – $T_3$ ($O_4$) Flank 1<br>$T_{17}$ ($O_4$) Loop 2 – $T_3$ ($N_3$) Flank 1 | 2.9<br>2.8 | Hydrogen bonds |
| | $A_8$ ($N_6$) Loop 1 – $A_{22}$ ($N_1$) Loop 3<br>$A_{22}$ ($N_6$) Loop 3 – $A_8$ ($N_7$) Loop 1 | 2.7<br>3.6 | Hydrogen bonds |
| **AT interactions** | | | |
| Strand | Nucleotides and atoms involved | Distance (Å) | Type of interaction |
| A | $A_{16}$($N_6$) Loop 2 – $T_{15}$($O_2$) Loop 2 | 3.1 | Hydrogen bond |
| **π–π stacking** | | | |
| Strand | Nucleotides involved | Distance (Å) | Type of interaction |
| A | $A_8$ Loop 1 – $A_{10}$ Loop 1 | - | π–π stacking |
| | $A_{10}$ Loop 1 – $C_7$ | - | π–π stacking |
| | $A_{16}$ Loop 2 – $T_{17}$ Loop 2 | - | π–π stacking |
| | $T_{29}$ Flank 2 – $A_{30}$ Flank 2 | - | π–π stacking |
| B | $C_7$ – $A_8$ Loop 1 | - | π–π stacking |
| | $C_{21}$ – $A_{22}$ Loop 3 | - | π–π stacking |
| **Interactions between nucleotides and the phosphate backbone** | | | |
| Strand | Nucleotides involved | Distance (Å) | Type of interaction |
| B | $A_{16}$($N_6$) Loop 2 – $A_{24}$ (OP1) Loop 3<br>$A_{16}$($N_6$) Loop 2 – $A_{24}$ (OP2) Loop 3 | 2.8<br>3.1 | Hydrogen bonds<br>($A_{16}$ is disordered) |

**Table S7.** Intermolecular interactions between strands A and B of the 4C i-motif based on the crystal structure.

$5'-T_1A_2T_3C_4C_5C_6C_7A_8C_9A_{10}C_{11}C_{12}C_{13}C_{14}T_{15}A_{16}T_{17}C_{18}C_{19}C_{20}C_{21}A_{22}C_{23}A_{24}C_{25}C_{26}C_{27}C_{28}T_{29}A_{30}T_{31}-3'$

Flank 1    Loop 1    Loop 2    Loop 3    Flank 2

| **Mismatched AA base pairs** | | | |
|---|---|---|---|
| **Strand A** nucleotides and atoms involved | **Strand B** nucleotides and atoms involved | Distance (Å) | Type of interaction |
| $A_2$ ($N_6$) Flank 1<br>$A_2$ ($N_1$) Flank 1 | $A_{24}$ ($N_7$) Loop 3<br>$A_{24}$ ($N_6$) Loop 3 | 2.8<br>2.7 | Hydrogen bonds |
| $A_{24}$ ($N_6$) Loop 3<br>$A_{24}$ ($N_7$) Loop 3 | $A_2$ ($N_1$) Flank 1<br>$A_2$ ($N_6$) Flank 1 | 3.0<br>3.1 | Hydrogen bonds |
| **AT base pairs** | | | |
| **Strand A** nucleotides and atoms involved | **Strand B** nucleotides and atoms involved | Distance (Å) | Type of interaction |
| $A_8$ ($N_6$) Loop 1<br>$A_8$ ($N_1$) Loop 1<br><br>$A_8$ ($N_6$) Loop 1<br>$A_8$ ($N_7$) Loop 1 | $T_{15}$ ($O_2$) Loop 2<br>$T_{15}$ ($N_3$) Loop 2<br><br>$T_{29}$ ($O_4$) Flank 2<br>$T_{29}$ ($N_3$) Flank 2 | 3.2<br>2.8<br><br>3.0<br>2.9 | **TAT triad**<br>Hydrogen bonds<br><br>Hoogsteen base pairing |
| $T_{15}$ ($O_4$) Loop 2<br>$T_{15}$ ($N_3$) Loop 2 | $A_{10}$ ($N_6$) Loop 1<br>$A_{10}$ ($N_1$) Loop 1 | 3.0<br>3.0 | Watson and Crick base pairing |
| **CC base pairs** | | | |
| **Strand A** nucleotides and atoms involved | **Strand B** nucleotides and atoms involved | Distance (Å) | Type of interaction |
| $C_9$ ($N_4$) Loop 1<br>$C_9$ ($N_3$) Loop 1<br>$C_9$ ($O_2$) Loop 1 | $C_{23}$($O_2$) Loop 3<br>$C_{23}$($N_3$) Loop 3<br>$C_{23}$ ($N_4$) Loop 3 | 2.9<br>2.8<br>2.9 | Hydrogen bonds |
| $C_{23}$($O_2$) Loop 3<br>$C_{23}$($N_3$) Loop 3<br>$C_{23}$ ($N_4$) Loop 3 | $C_9$ ($N_4$) Loop 1<br>$C_9$ ($N_3$) Loop 1<br>$C_9$ ($O_2$) Loop 1 | 2.9<br>2.8<br>2.9 | Hydrogen bonds |
| **π–π stacking** | | | |
| **Strand A** nucleotides and atoms involved | **Strand B** nucleotides and atoms involved | Distance (Å) | Type of interaction |
| $A_{24}$ Loop 3 | $T_1$ Flank 1 | - | π–π stacking |
| $T_1$ Flank 1 | $A_{24}$ Loop 3 | - | π–π stacking |
| $A_2$ Flank 1 | $C_9$ Loop 1 | - | π–π stacking |
| $T_{15}$ Loop 2 | $T_{31}$ Flank 2 | - | π–π stacking<br>($T_{31}$ is disordered) |
| $A_{16}$ Loop 2 | $A_{22}$ Loop 3 | - | π–π stacking |
| $A_{22}$ Loop 3 | $T_{15}$ Loop 2 | - | π–π stacking |
| $C_9$ Loop 1 | $A_2$ Flank 1 | - | π–π stacking |

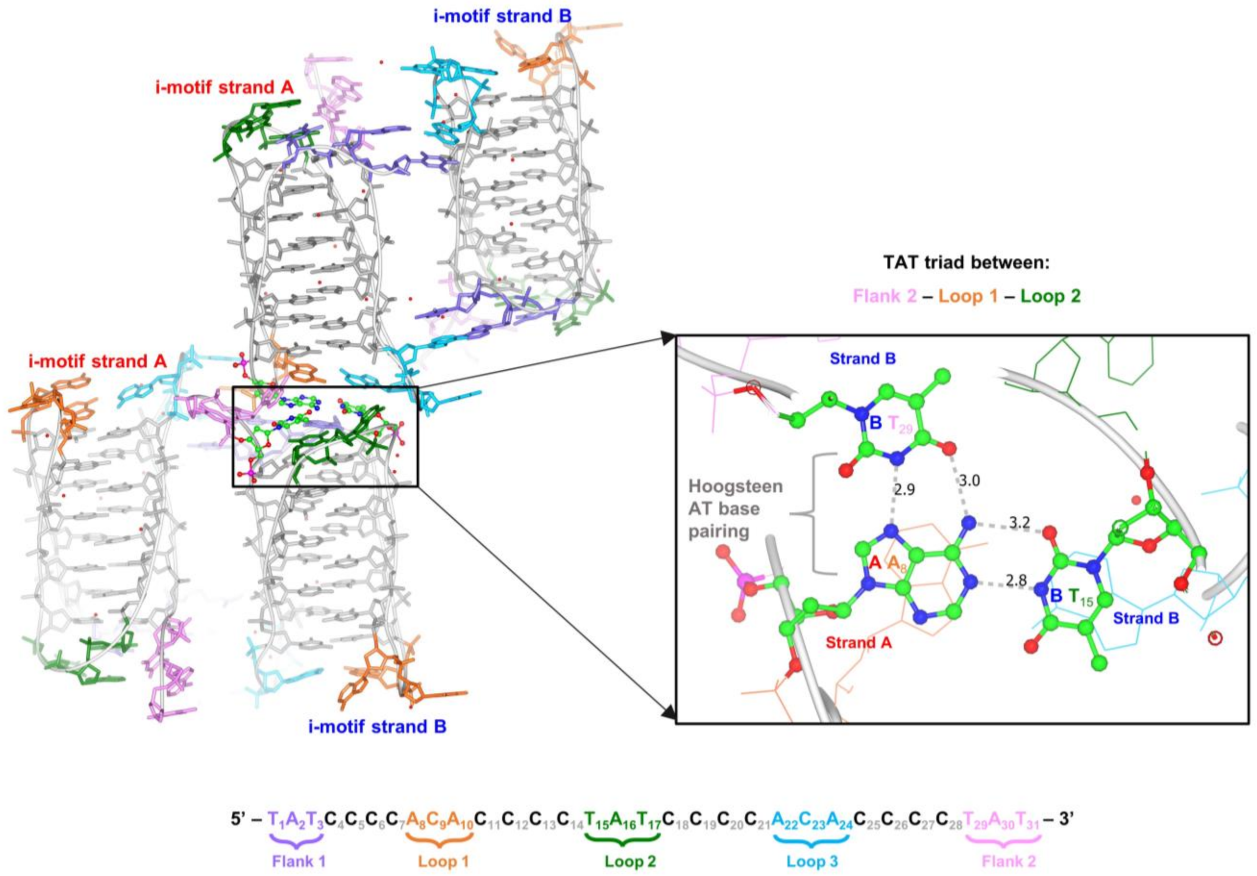


**Figure S12.** Intramolecular 4C i-motifs A and B forming a TAT triad.

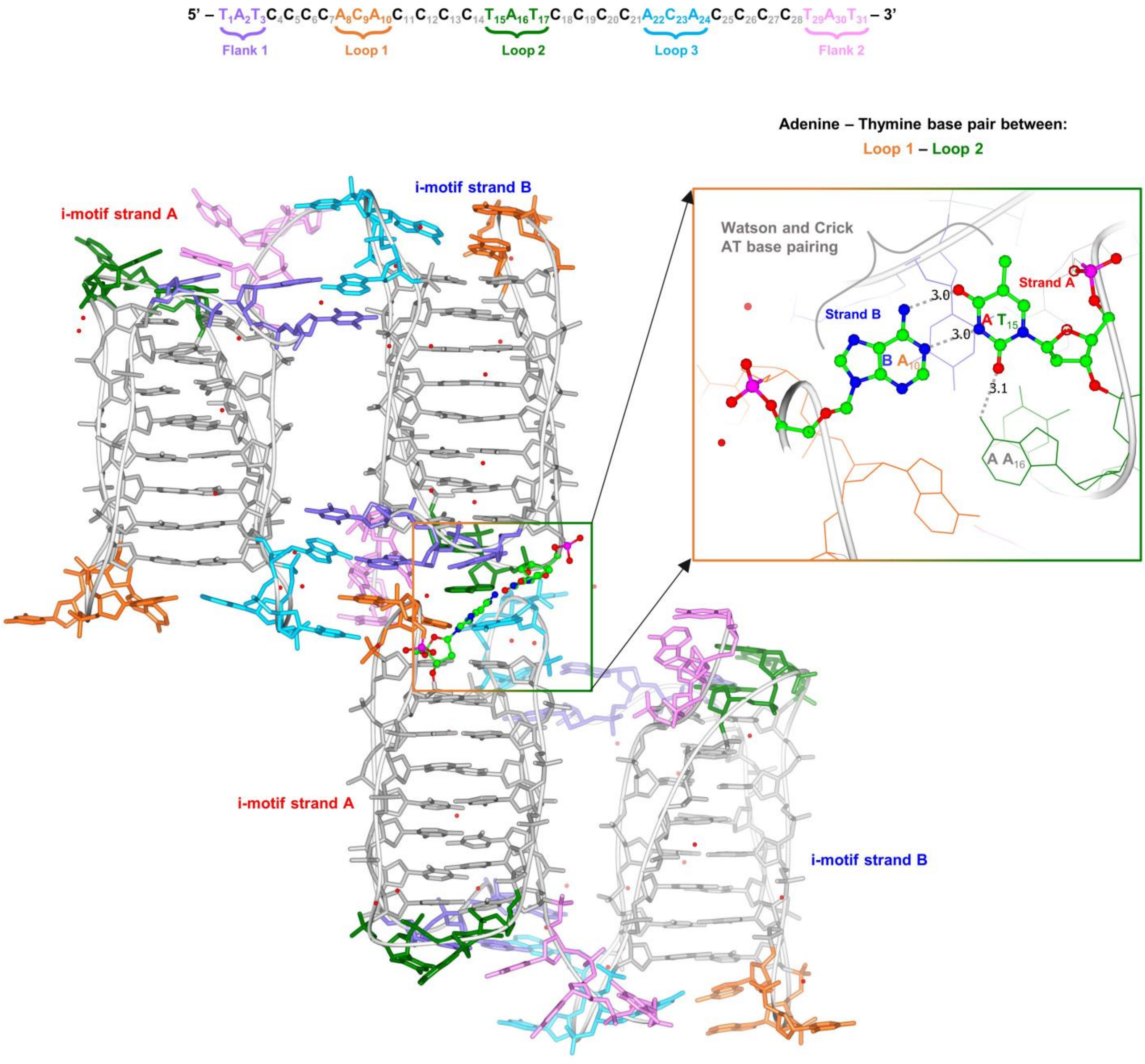


**Figure S13.** Intramolecular 4C i-motifs A and B forming AT base pairs.

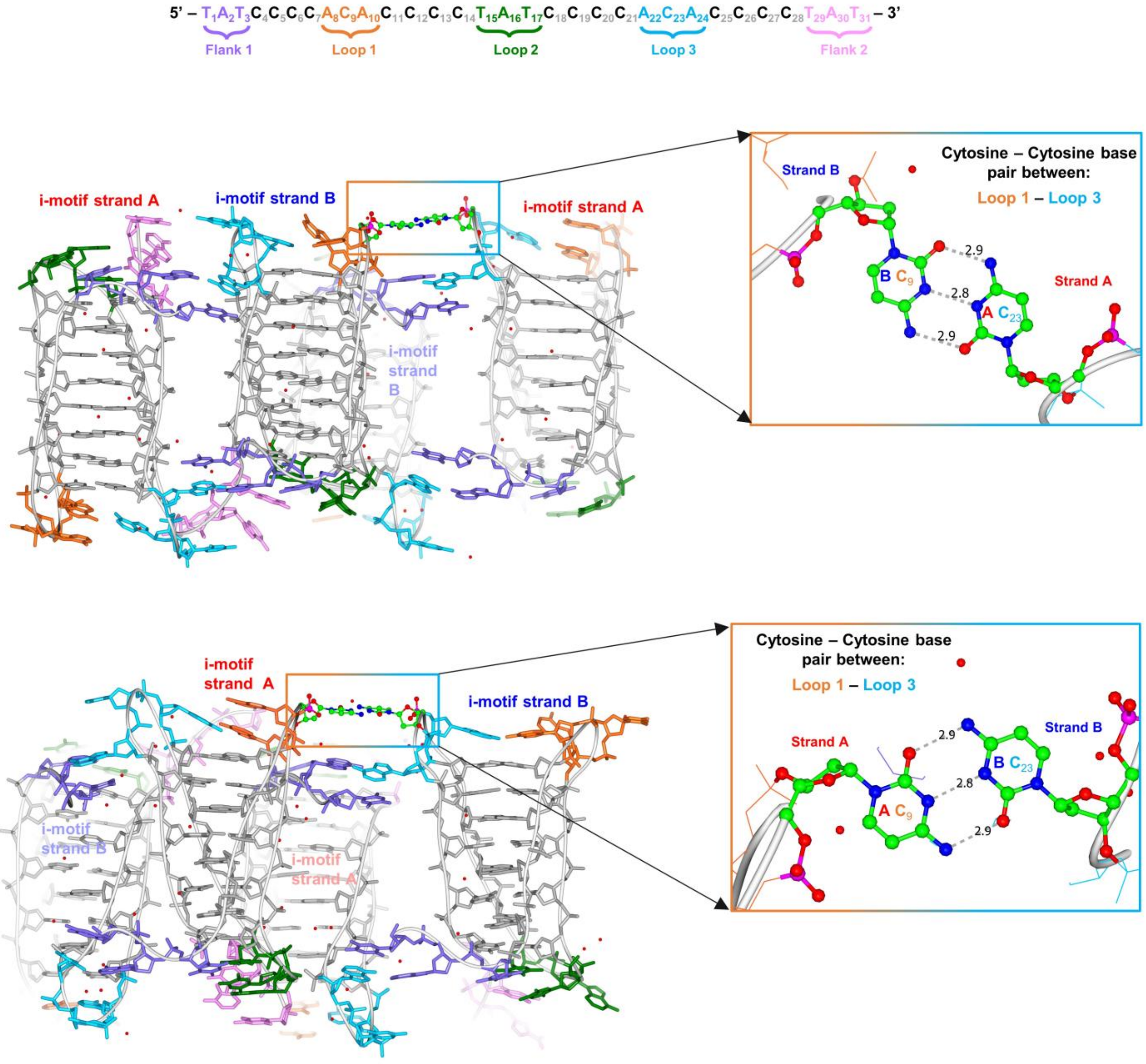


**Figure S14.** Intramolecular 4C i-motifs A and B forming CC base pairs.

**Table S8.** Nucleotide backbone torsion angles and sugar puckers in the two strands of the 4C i – motif calculated by the 3DNA.[2] $\Delta_{A-B}$ = A-B , $\Delta_{A-B}$*= absolute angle difference between strand A and B within the range of 0° to 180°.

| Base | Strand | Alpha | Beta | Gamma | Delta | Epsilon | Zeta | Chi | Orientation | Puckering |
|---|---|---|---|---|---|---|---|---|---|---|
| T1 | A | --- | --- | 70.8 | 149.2 | -98.6 | 78.9 | -133 | anti | C2' - endo |
| | B | --- | --- | -40.3 | 159 | -171.7 | 145.7 | -142.6 | anti | C3' - exo |
| | $\Delta_{A-B}$ | --- | --- | 111.1 | -9.8 | 73.1 | -66.8 | 9.6 | | |
| | $\Delta_{A-B}$* | --- | --- | 111.1 | 9.8 | 73.1 | 66.8 | 9.6 | | |
| A2 | A | -161.3 | -143.1 | 68.4 | 102.3 | -85.1 | 178.5 | -77.4 | syn | C2' - exo |
| | B | -58.5 | 136.6 | 47.8 | 112.5 | -104.1 | -152.3 | -71.3 | syn | C2' - exo |
| | $\Delta_{A-B}$ | -102.8 | -279.7 | 20.6 | -10.2 | 19 | 330.8 | -6.1 | | |
| | $\Delta_{A-B}$* | 102.8 | 80.3 | 20.6 | 10.2 | 19 | 29.2 | 6.1 | | |
| T3 | A | -35.3 | -166.4 | 41.6 | 145.8 | -129 | -80.1 | -146.3 | anti | C3' - exo |
| | B | -24.3 | -163.9 | 44.2 | 148.8 | -118.9 | -65.3 | -158.9 | anti | C3' - exo |
| | $\Delta_{A-B}$ | -11 | -2.5 | -2.6 | -3 | -10.1 | -14.8 | 12.6 | | |
| | $\Delta_{A-B}$* | 11 | 2.5 | 2.6 | 3 | 10.1 | 14.8 | 12.6 | | |
| C4 | A | -58.5 | 169.6 | 52.7 | 128.3 | -126 | -121.6 | -106.7 | anti | C2' - endo |
| | B | -60.4 | 166.6 | 37.5 | 94.7 | -166 | -109.4 | -120.7 | anti | C4' - exo |
| | $\Delta_{A-B}$ | 1.9 | 3 | 15.2 | 33.6 | 40 | -12.2 | 14 | | |
| | $\Delta_{A-B}$* | 1.9 | 3 | 15.2 | 33.6 | 40 | 12.2 | 14 | | |
| C5 | A | 67.8 | -158.1 | -112.6 | 130.7 | -161.5 | -111.8 | -92.1 | anti | C2' - endo |
| | B | 142.1 | -162.8 | -151.9 | 116.2 | -131 | -126.5 | -124.4 | anti | C1' - exo |
| | $\Delta_{A-B}$ | -74.3 | 4.7 | 39.3 | 14.5 | -30.5 | 14.7 | 32.3 | | |
| | $\Delta_{A-B}$* | 74.3 | 4.7 | 39.3 | 14.5 | 30.5 | 14.7 | 32.3 | | |
| C6 | A | 174.4 | 174.1 | 163.5 | 84.9 | 67.6 | 75.3 | -124.1 | anti | C4' - exo |
| | B | 94.6 | -150.6 | -130.6 | 106.8 | -149.6 | -41 | -127.5 | anti | C1' - exo |
| | $\Delta_{A-B}$ | 79.8 | 324.7 | 294.1 | -21.9 | 217.2 | 116.3 | 3.4 | | |
| | $\Delta_{A-B}$* | 79.8 | 35.3 | 65.9 | 21.9 | 142.8 | 116.3 | 3.4 | | |
| C7 | A | 97.1 | 163.9 | 174.5 | 141.1 | -103.2 | -119.8 | -103.7 | anti | C2' - endo |
| | B | -142.1 | 84 | -171.4 | 136.3 | -127.8 | -139.5 | -122 | anti | C2' - endo |
| | $\Delta_{A-B}$ | 239.2 | 79.9 | 345.9 | 4.8 | 24.6 | 19.7 | 18.3 | | |
| | $\Delta_{A-B}$* | 120.8 | 79.9 | 14.1 | 4.8 | 24.6 | 19.7 | 18.3 | | |
| A8 | A | 7 | -114.1 | -90.2 | 164.9 | -149.3 | -178.7 | -98.8 | anti | C3' - exo |
| | B | 97.2 | -155 | -160.9 | 142 | -96.8 | 150.6 | 64.8 | syn | C2' - endo |
| | $\Delta_{A-B}$ | -90.2 | 40.9 | 70.7 | 22.9 | -52.5 | -329.3 | -163.6 | | |
| | $\Delta_{A-B}$* | 90.2 | 40.9 | 70.7 | 22.9 | 52.5 | 30.7 | 163.6 | | |
| C9 | A | 62.1 | 174.7 | 53.6 | 94.8 | -139.1 | 90 | -139.2 | anti | C3' - endo |
| | B | 72.5 | 150.6 | 40.4 | 89.5 | -174.9 | 109.6 | -156.9 | anti | C3' - endo |
| | $\Delta_{A-B}$ | -10.4 | 24.1 | 13.2 | 5.3 | 35.8 | -19.6 | 17.7 | | |
| | $\Delta_{A-B}$* | 10.4 | 24.1 | 13.2 | 5.3 | 35.8 | 19.6 | 17.7 | | |
| A10 | A | -91.6 | -138.4 | 47.5 | 90.5 | -173.7 | -100.8 | -114.5 | anti | C4' - exo |
| | B | -62 | 143.5 | 42.6 | 110.5 | -142.8 | 168.2 | -134.3 | anti | C1' - exo |
| | $\Delta_{A-B}$ | -29.6 | -281.9 | 4.9 | -20 | -30.9 | -269 | 19.8 | | |
| | $\Delta_{A-B}$* | 29.6 | 78.1 | 4.9 | 20 | 30.9 | 91 | 19.8 | | |
| C11 | A | -49.5 | 173.1 | 55.6 | 90.1 | -152.6 | -92.5 | -128.6 | anti | C3' - endo |
| | B | -47.2 | 170.9 | 46 | 91.6 | -122.5 | -114 | -140.5 | anti | C3' - endo |
| | $\Delta_{A-B}$ | -2.3 | 2.2 | 9.6 | -1.5 | -30.1 | 21.5 | 11.9 | | |
| | $\Delta_{A-B}$* | 2.3 | 2.2 | 9.6 | 1.5 | 30.1 | 21.5 | 11.9 | | |
| C12 | A | -49.2 | 171.5 | 54.2 | 85.7 | -172.8 | -73.7 | -126.4 | anti | C3' - endo |
| | B | -27.5 | 148.4 | 39.1 | 80.2 | -160.8 | -101.7 | -124.9 | anti | C3' - endo |
| | $\Delta_{A-B}$ | -21.7 | 23.1 | 15.1 | 5.5 | -12 | 28 | -1.5 | | |
| | $\Delta_{A-B}$* | 21.7 | 23.1 | 15.1 | 5.5 | 12 | 28 | 1.5 | | |
| C13 | A | -86.4 | -166.6 | 74.6 | 81.9 | -153 | -90.5 | -122.9 | anti | C3' - endo |
| | B | -42.7 | 158.1 | 59.1 | 81.7 | -157.6 | -90 | -132.9 | anti | C3' - endo |
| | $\Delta_{A-B}$ | -43.7 | -324.7 | 15.5 | 0.2 | 4.6 | -0.5 | 10 | | |
| | $\Delta_{A-B}$* | 43.7 | 35.3 | 15.5 | 0.2 | 4.6 | 0.5 | 10 | | |
| C14 | A | -47.3 | 167 | 57.5 | 79.4 | -145.8 | -15.9 | -120.2 | anti | C3' - endo |
| | B | 146.4 | -172.8 | -158.5 | 90.7 | -113.6 | -56 | -125.2 | anti | C3' - endo |

| | | | | | | | | | | |
|---|---|---|---|---|---|---|---|---|---|---|
| | $\Delta_{A-B}$ | -193.7 | 339.8 | 216 | -11.3 | -32.2 | 40.1 | 5 | | |
| | $\Delta_{A-B}$* | 166.3 | 20.2 | 144 | 11.3 | 32.2 | 40.1 | 5 | | |
| T15 | A | 177.9 | 117.4 | 77.6 | 121.1 | -106.8 | -143.5 | -175.4 | anti | C1' - exo |
| | B | 153 | -166.1 | 77.1 | 145.5 | -147.4 | -160.1 | -142.1 | anti | C2' - endo |
| | $\Delta_{A-B}$ | 24.9 | 283.5 | 0.5 | -24.4 | 40.6 | 16.6 | -33.3 | | |
| | $\Delta_{A-B}$* | 24.9 | 76.5 | 0.5 | 24.4 | 40.6 | 16.6 | 33.3 | | |
| A16 | A | -173.2 | -160.6 | 77.7 | 89.8 | -137.6 | -66.6 | -174.5 | anti | C3' - endo |
| | B | 58.9 | 121.4 | 168.4 | 147.7 | 19.4 | -173.5 | 69.2 | syn | C3' - exo |
| | $\Delta_{A-B}$ | -232.1 | -282 | -90.7 | -57.9 | -157 | 106.9 | -243.7 | | |
| | $\Delta_{A-B}$* | 127.9 | 78 | 90.7 | 57.9 | 157 | 106.9 | 116.3 | | |
| T17 | A | -65.5 | -177.2 | 49.5 | 86.8 | -133.9 | -87.5 | -154.2 | anti | C3' - endo |
| | B | -29.6 | 136.9 | 11.5 | 85.1 | -140.2 | -89.6 | -144.1 | anti | C3' - endo |
| | $\Delta_{A-B}$ | -35.9 | -314.1 | 38 | 1.7 | 6.3 | 2.1 | -10.1 | | |
| | $\Delta_{A-B}$* | 35.9 | 45.9 | 38 | 1.7 | 6.3 | 2.1 | 10.1 | | |
| C18 | A | -55 | -179.7 | 55.5 | 89.1 | -167.6 | -78.7 | -125.2 | anti | C3' - endo |
| | B | -45.2 | 178 | 52.6 | 91.8 | -163.2 | -91.6 | -126 | anti | C3' - endo |
| | $\Delta_{A-B}$ | -9.8 | -357.7 | 2.9 | -2.7 | -4.4 | 12.9 | 0.8 | | |
| | $\Delta_{A-B}$* | 9.8 | 2.3 | 2.9 | 2.7 | 4.4 | 12.9 | 0.8 | | |
| C19 | A | -68.8 | -167.5 | 61.7 | 89.9 | -162.5 | -89.6 | -118.7 | anti | C3' - endo |
| | B | -53.9 | 179.9 | 51 | 87.3 | -170.3 | -80.4 | -115.1 | anti | C3' - endo |
| | $\Delta_{A-B}$ | -14.9 | -347.4 | 10.7 | 2.6 | 7.8 | -9.2 | -3.6 | | |
| | $\Delta_{A-B}$* | 14.9 | 12.6 | 10.7 | 2.6 | 7.8 | 9.2 | 3.6 | | |
| C20 | A | -63.7 | 170.7 | 69 | 86.5 | -158.3 | -84.9 | -129 | anti | C3' - endo |
| | B | -64.5 | 179.8 | 66.1 | 85.5 | -149 | -111.2 | -120.4 | anti | C3' - endo |
| | $\Delta_{A-B}$ | 0.8 | -9.1 | 2.9 | 1 | -9.3 | 26.3 | -8.6 | | |
| | $\Delta_{A-B}$* | 0.8 | 9.1 | 2.9 | 1 | 9.3 | 26.3 | 8.6 | | |
| C21 | A | -44.4 | -171.3 | 49.7 | 95.7 | -144.1 | -124.1 | -103.3 | anti | C4' - exo |
| | B | 79.3 | -141.6 | -85.7 | 162.1 | -115.9 | -127.2 | -97.9 | anti | C3' - exo |
| | $\Delta_{A-B}$ | -123.7 | -29.7 | 135.4 | -66.4 | -28.2 | 3.1 | -5.4 | | |
| | $\Delta_{A-B}$* | 123.7 | 29.7 | 135.4 | 66.4 | 28.2 | 3.1 | 5.4 | | |
| A22 | A | -42.9 | 166.6 | 48.4 | 155.7 | -98.1 | 96.3 | -55.1 | syn | C3' - exo |
| | B | 81.4 | -138.9 | -158.5 | 144.8 | -93.5 | 134.7 | -92.8 | anti | C2' - endo |
| | $\Delta_{A-B}$ | -124.3 | 305.5 | 206.9 | 10.9 | -4.6 | -38.4 | 37.7 | | |
| | $\Delta_{A-B}$* | 124.3 | 54.5 | 153.1 | 10.9 | 4.6 | 38.4 | 37.7 | | |
| C23 | A | 73 | 177.7 | 28.4 | 89.3 | -153.7 | -90.5 | -147.2 | anti | C3' - endo |
| | B | 39.3 | 165.1 | 74.9 | 140.2 | -122.7 | -40.9 | -142 | anti | C2' - endo |
| | $\Delta_{A-B}$ | 33.7 | 12.6 | -46.5 | -50.9 | -31 | -49.6 | -5.2 | | |
| | $\Delta_{A-B}$* | 33.7 | 12.6 | 46.5 | 50.9 | 31 | 49.6 | 5.2 | | |
| A24 | A | -49.1 | -144.5 | 40.8 | 145 | -103.7 | -67.5 | -90.8 | anti | C2' - endo |
| | B | -70.6 | -123.2 | -70.1 | 165.3 | -109.5 | -92.6 | -66.9 | syn | C3' - exo |
| | $\Delta_{A-B}$ | 21.5 | -21.3 | 110.9 | -20.3 | 5.8 | 25.1 | -23.9 | | |
| | $\Delta_{A-B}$* | 21.5 | 21.3 | 110.9 | 20.3 | 5.8 | 25.1 | 23.9 | | |
| C25 | A | -62.3 | 174.4 | 173.3 | 138.4 | -153.1 | -56.3 | -137.1 | anti | C2' - endo |
| | B | 62.5 | 159.2 | 53.7 | 141.2 | -153.6 | -66.7 | -108.5 | anti | C2' - endo |
| | $\Delta_{A-B}$ | -124.8 | 15.2 | 119.6 | -2.8 | 0.5 | 10.4 | -28.6 | | |
| | $\Delta_{A-B}$* | 124.8 | 15.2 | 119.6 | 2.8 | 0.5 | 10.4 | 28.6 | | |
| C26 | A | 154.4 | 142 | -177.1 | 118.3 | -156.9 | -72.7 | -109.6 | anti | C1' - exo |
| | B | 167.7 | 135.1 | 176.9 | 120.5 | -161.6 | -73.8 | -106.9 | anti | C1' - exo |
| | $\Delta_{A-B}$ | -13.3 | 6.9 | -354 | -2.2 | 4.7 | 1.1 | -2.7 | | |
| | $\Delta_{A-B}$* | 13.3 | 6.9 | 6 | 2.2 | 4.7 | 1.1 | 2.7 | | |
| C27 | A | -157.5 | 122.3 | 171 | 128.2 | -150.2 | -75.8 | -113.5 | anti | C2' - endo |
| | B | 179.1 | 138.3 | 170.1 | 91 | -135.7 | -72.6 | -116.7 | anti | C4' - exo |
| | $\Delta_{A-B}$ | -336.6 | -16 | 0.9 | 37.2 | -14.5 | -3.2 | 3.2 | | |
| | $\Delta_{A-B}$* | 23.4 | 16 | 0.9 | 37.2 | 14.5 | 3.2 | 3.2 | | |
| C28 | A | -175.4 | 134.9 | 169.9 | 112.7 | -88.6 | 134.9 | -108.2 | anti | C1' - exo |
| | B | -104.7 | 98.2 | 165.3 | 136.3 | -113.2 | -68.3 | -139 | anti | C2' - endo |
| | $\Delta_{A-B}$ | -70.7 | 36.7 | 4.6 | -23.6 | 24.6 | 203.2 | 30.8 | | |
| | $\Delta_{A-B}$* | 70.7 | 36.7 | 4.6 | 23.6 | 24.6 | 156.8 | 30.8 | | |
| T29 | A | 3.1 | 123.3 | 9.1 | 150.5 | -167.2 | -136.3 | -107.8 | anti | C2' - endo |
| | B | -56.5 | -128.1 | -62.1 | 136.2 | -151.5 | -118.4 | 88 | syn | C2' - endo |
| | $\Delta_{A-B}$ | 59.6 | 251.4 | 71.2 | 14.3 | -15.7 | -17.9 | -195.8 | | |
| | $\Delta_{A-B}$* | 59.6 | 108.6 | 71.2 | 14.3 | 15.7 | 17.9 | 164.2 | | |

| | | | | | | | | | | |
|---|---|---|---|---|---|---|---|---|---|---|
| A30 | A | 16 | 129.3 | 16.3 | 140.9 | -179 | 163 | -86 | syn | C2' - endo |
| | B | 41.7 | -122.9 | -100.4 | 70.9 | -149.8 | 171.6 | -159.7 | anti | C3' - endo |
| | $\Delta_{A-B}$ | -25.7 | 252.2 | 116.7 | 70 | -29.2 | -8.6 | 73.7 | | |
| | $\Delta_{A-B}$* | 25.7 | 107.8 | 116.7 | 70 | 29.2 | 8.6 | 73.7 | | |
| T31 | A | 15.5 | 113.5 | 8.3 | 89.9 | --- | --- | -90.9 | anti | C3' - endo |
| | B | 36.8 | 122.7 | -8.7 | 150.1 | --- | --- | -151.9 | anti | C2' - endo |
| | $\Delta_{A-B}$ | -21.3 | -9.2 | 17 | -60.2 | --- | --- | 61 | | |
| | $\Delta_{A-B}$* | 21.3 | 9.2 | 17 | 60.2 | --- | --- | 61 | | |

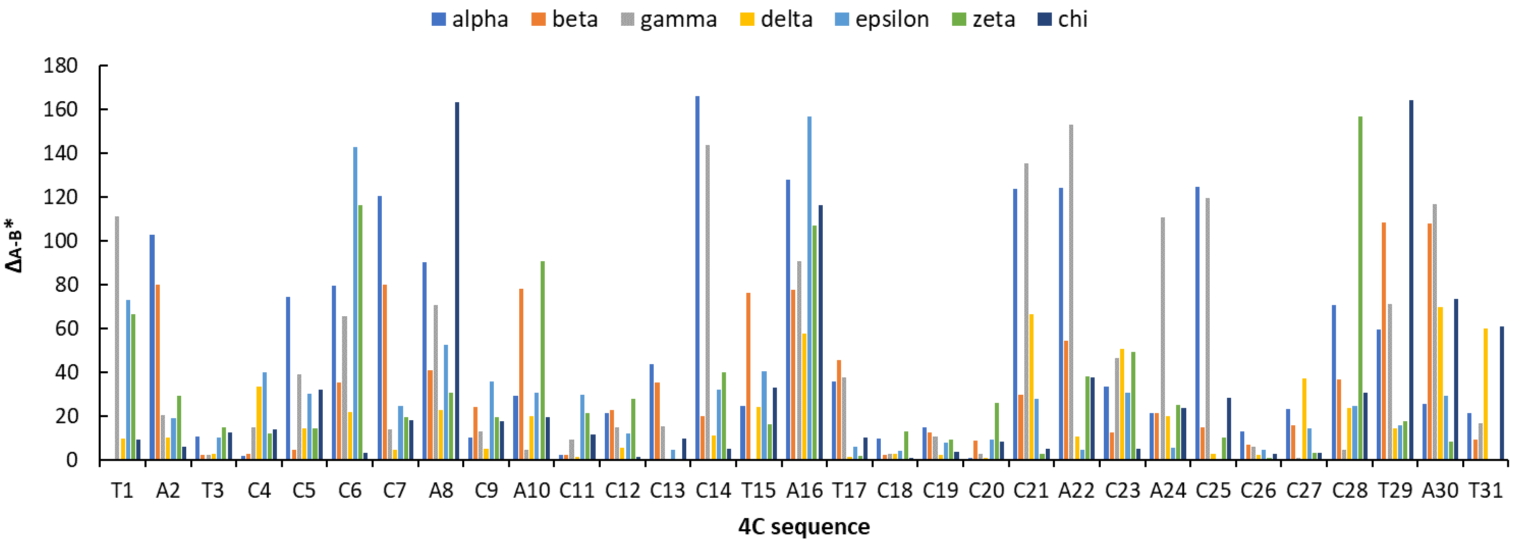


**Figure S15.** Comparison of the torsion angles and sugar pucker between strand A and strand B. The smallest absolute difference between strand A and B within the range of 0° to 180° ($\Delta_{A-B}$* ) is shown for each nucleotide of the 4C sequence.

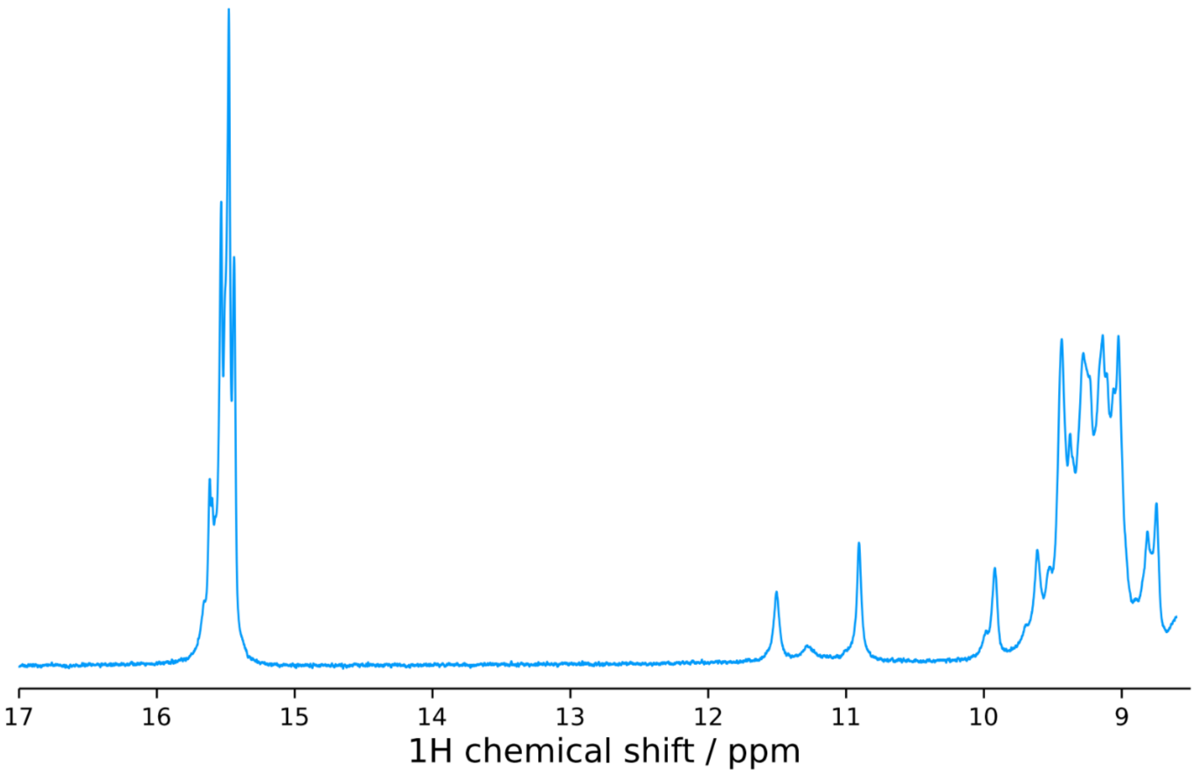


**Figure S16:** $^{1}$H-1D NMR of 4C 0.7 mM, 9.1 mM NaCaco, pH 5.5, 91 mM KCl, 17% $D_2O$.

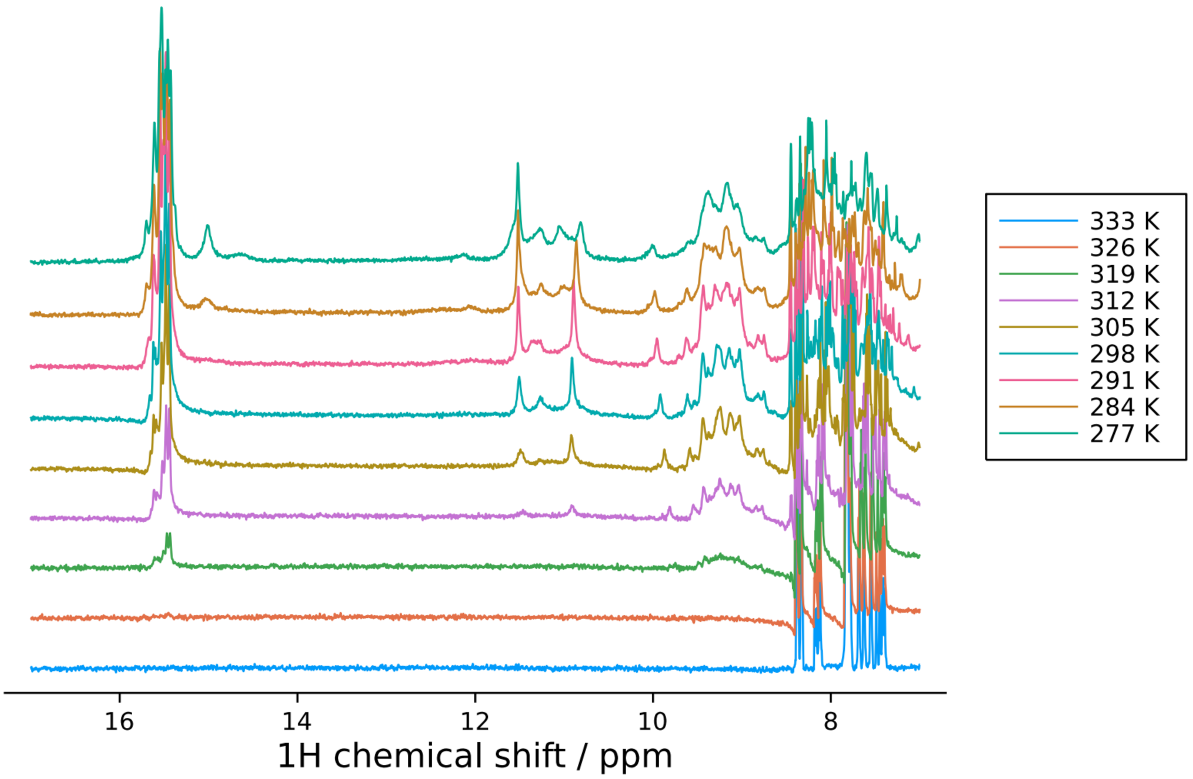


**Figure S17:** Annealing $^{1}$H-1D NMR of 4C 0.7 mM, 9.1 mM NaCaco, pH 5.5, 91 mM KCl, 17% $D_2O$ between 333 and 277 K.

**Supplementary Information for enhanced sampling molecular dynamics simulations**

The initial ILPR i-motif crystal structure consists of two near-identical motifs packed into a dimer as a result of crystallisation. Given the interactions between the sequences observed in the crystal structure originate from the flanking sequence, we looked at sequence 4C (TATCCCCACACCCCTATCCCCACACCCCTAT) and also an analogue with one base missing at the 3'-end of the flanking sequence, 4Cdel (TATCCCCACACCCCTATCCCCACACCCCTA).

The structures were prepared using the *tleap* module of the AmberTools20 package.[3] With regards to force-field parameters, a hybrid model, combining both bsc1 and OL15 modifications were used.[4,5] As with all AMBER force-fields, these are additive and can be combined. Bsc1 is currently the best descriptor of alpha/gamma torsion angles, while OL15 best describes ε/**ζ** dihedrals and non-bonded interactions of bases. Further to these force-field parameters for protonated cytosine were taken from the AMBER force-field library: 'all_prot_nucleic10.lib'. In the case of the protonated structures, two of the four strands were protonated at the N3 position ($C_{4-7}$ and $C_{11-14}$), while the parameters mentioned previously apply to the remaining nucleotides.

Sufficient $K^+$ was added to the systems to neutralise the negatively charged phosphate backbones, as well as the C+ atoms in the case of the protonated systems. Then excess $K^+$ and $Cl^-$ atoms were added to simulate a 0.15 mol $L^{-1}$ salt concentration. The systems were solvated using the TIP3P water model in an orthorhombic box, with all solute atoms at a minimum distance of 12 Å from the edge of the solvation box.

Each system was minimised and equilibrated under NVT conditions for 5 ns at 1 atm. The temperature was set to 300K using a timestep of 4 fs, all rigid bonds, a cut-off of 9 Å and particle mesh Ewald summation (PME) switched on for long-range electrostatics. The nucleic acid was initially fixed during equilibration, but these restraints were gradually removed, while the ions and solvent molecules were allowed to move. The heavy atoms of the nucleic acid were constrained by a spring constant set at 1 kcal $mol^{-1}$ $Å^{-2}$. The production simulations were run in the NVT ensemble using a Langevin thermostat with a damping of 1.0 ps−1 and hydrogen mass repartitioning scheme to achieve time steps of 4 fs. The AdaptiveBandit algorithm is implemented here,[6] and after each short run (50 ns) the algorithm is given a decision to select a pre-sampled conformation from which to restart the next short simulation - thus the objective is to avoid redundant sampling (i.e. visiting states which have already been sampled). The decision that is made aims to maximise a reward function. Here the reward function was defined to be proportional to relative phosphate-phosphate distances throughout the structure.

The decision made each time aims to maximise the conformational space explored, using phosphate-phosphate distances as a metric.

The i-motif loops represent distinct sampling problems in that the core structure, stabilised by the hemi-protonated cytosine pairs, is largely stable with minor fluctuations. However, the loop regions are highly dynamic, able to sample multiple conformations with varying degrees of stability. The phosphate-phosphate distances from the structure were used as the Adaptive Bandit metric for the first 20 µs of simulation, after this only phosphate-phosphate distances in loop regions (8-10, 15-17, 22-24) were used as metrics. The loop-only sampling systems were run for a total of 30 µs. Thus, each system was run for a total sampling time of atleast 50 µs.

As previously mentioned, i-motif loops represent a distinct sampling problem and conventional molecular dynamics simulations are not able to fully explore the conformational space in which it exists. This is overcome by use of adaptive sampling algorithms such as Adaptive Bandit, however the output from these many short simulations cannot be analysed directly therefore we need to integrate these simulations into Markov state models (MSMs). MSMs allow the integration of multiple simulation trajectories into a single model of the conformational landscape that contains both kinetic and thermodynamic information as well as maintain structural detail. MSMs are built on inter-state transitions; all of these states can be compiled and used to create a single model.

Here we used the PyEMMA package to create our models.[7] The general workflow for creating a Markov goes as follows. First, we use Time-lagged Independent Component Analysis (tICA) to reduce the dimensionality of the input feature. This differs from Principal Component Analysis in that it expands this reduced representation into the time domain. The data from tICA is then projected onto the first two (i.e. two slowest) ICs at a selected lag time and then clustered using the k-means algorithm. The number of clusters chosen represents the number of clusters at which the VAMP2 scoring function no longer increases. This allows the numerous conformations to be mapped onto a set of discrete states. This then allows MSMs to be created that describe transitions that occur between any two clusters. The MSM calculates these transitions between states at a given lag time. It should be noted that this lag time is independent from that used in tICA. Choice of MSM lag time is key in that it must be sufficiently long to ensure Markovian dynamics (i.e. the system is memory-less and each transition isn't dependent on the previous state) but short enough to still be able to resolve the dynamics of the system. In order to choose this lag time we choose the smallest possible lag time at which the underlying processes (represented in an implied-timescale plot) show convergence. This model can then be validated using the Chapman-Kolmogorov test, which tests the ability of the model at longer time scales. The original space can then be discretized into the longest-living metastable states according to their eigenvectors. Then, the most stable structure is then extracted from each metastable state, as well as a distribution of 50 structures with an associated trajectory file to display transition between all 50 structures in a particular state. Finally the PCCA++ algorithm is used to coarse-grain the conformational space of the metastable states and then approximate the stationary distribution of the data as well as approximating the relative free energies.

A total of 6 Markov state models were created to describe different aspects of the dynamics of the i-motif structure, details of which are displayed in Table S9.

A combination of $\alpha/\gamma$ angles and the distance between N3 atoms of the stem cysteines were used as features to describe the central stem, while a combination of $\alpha/\gamma$ angles and $\chi$ angles were used as features to describe the dynamics of the loops.

| | **Features** | **tICA lag** | **N° dimensions** | **Clusters** | **MSM lag** | **N° states** |
|---|---|---|---|---|---|---|
| **4C stem** | $\alpha/\gamma$ dihedral + N3 distance of stem bases | 450 | 3 | 200 | 60 | 5 |
| **4C Loop 1 and 3** | $\alpha/\gamma$ dihedral + $\chi$ | 30 | 3 | 200 | 10 | 5 |
| **4C Loop 2** | $\alpha/\gamma$ dihedral + $\chi$ | 30 | 3 | 200 | 20 | 4 |
| **4Cdel stem** | $\alpha/\gamma$ dihedral + N3 distance of stem bases | 400 | 5 | 200 | 60 | 3 |
| **4Cdel Loop 1 and 3** | $\alpha/\gamma$ dihedral + $\chi$ | 40 | 3 | 200 | 20 | 6 |
| **4Cdel Loop 2** | $\alpha/\gamma$ dihedral + $\chi$ | 40 | 3 | 200 | 30 | 5 |

**Table S9.** Description of key parameters for creating Markov State Models for sequences 4C and 4Cdel

In the ILPR i-motif structure there are three loops, 1 ($A_8C_9A_{10}$) and 3 ($A_{22}C_{23}A_{24}$) are distal to the 5' and 3' ends, while loop 2 ($T_{15}A_{16}T_{17}$) lies in the middle at the sequence and is flanked in space by both the 5' and 3' prime ends. 4Cdel, loop 1 consists of $C_9$ protruding away from the centre of the structure and into the solvent, while $A_{10}$ intercalates between $A_8$ and $C_7$, interacting with both via $\pi$-stacking, as well as forming a non-canonical base pair with $A_{22}$. In loop 3, $A_{22}$ is pointed towards the centre of the structure and caps $C_{21}$, while $C_{23}$ and $C_{24}$ protrude into the solvent (as a result of crystal packing) with their faces around 6.5 Å apart. In loop 2, $T_{17}$ interacts with $T_3$ via two hydrogen bonds formed between (O4 and N3), this pair forms in parallel to the C·C$^+$ pair above and stabilises these via $\pi$-$\pi$ interactions. $T_{17}$ itself is stabilised via $\pi$-stacking with $T_{16}$ below. $T_{15}$ forms no obvious actions and looks to shield the internal structure of the i-motif from solvent. The 4C structure is near-identical apart from $A_{10}$, which instead of intercalating between $A_8$ and $C_7$, is flipped outward into the solvent, $A_8$ instead, forms the non-canonical base pair with $A_{22}$. This is also the case with $A_{16}$, which protrudes into the solvent instead of stacking below $T_{17,}$ which itself is instead stabilised from below by $T_{29}$.

In order to assess the individual components that contribute to structural stability within the i-motif, MSMs were created with features extracted from only loop regions (bases 8-10, 15-17 & 22-24) and then with features from stem regions of the structure (bases 4-7, 11-14, 18-21). As expected, the loop regions are far more flexible and the majority of dynamic motion within the structure can be attributed to these regions. This is borne out in the fact that the stem regions required significantly larger tICA (40 ns) and MSM lag times, than the models of the loop systems, to fully describe the conformational landscape. This is also displayed by the findings that the stem only exists in a two-state system, displayed by a double-well potential (Figure S18/19). These two states have an RMSD < 1.2 Å, whereas the conformational landscapes are split into many more metastable states with significantly higher RMSD values. Although not a perfect comparison, this agrees with similar work on G-quadruplex structures using similar force-fields, as such validating the fact that these are suitable to study i-motif stem structures and C·C$^+$ pairs.

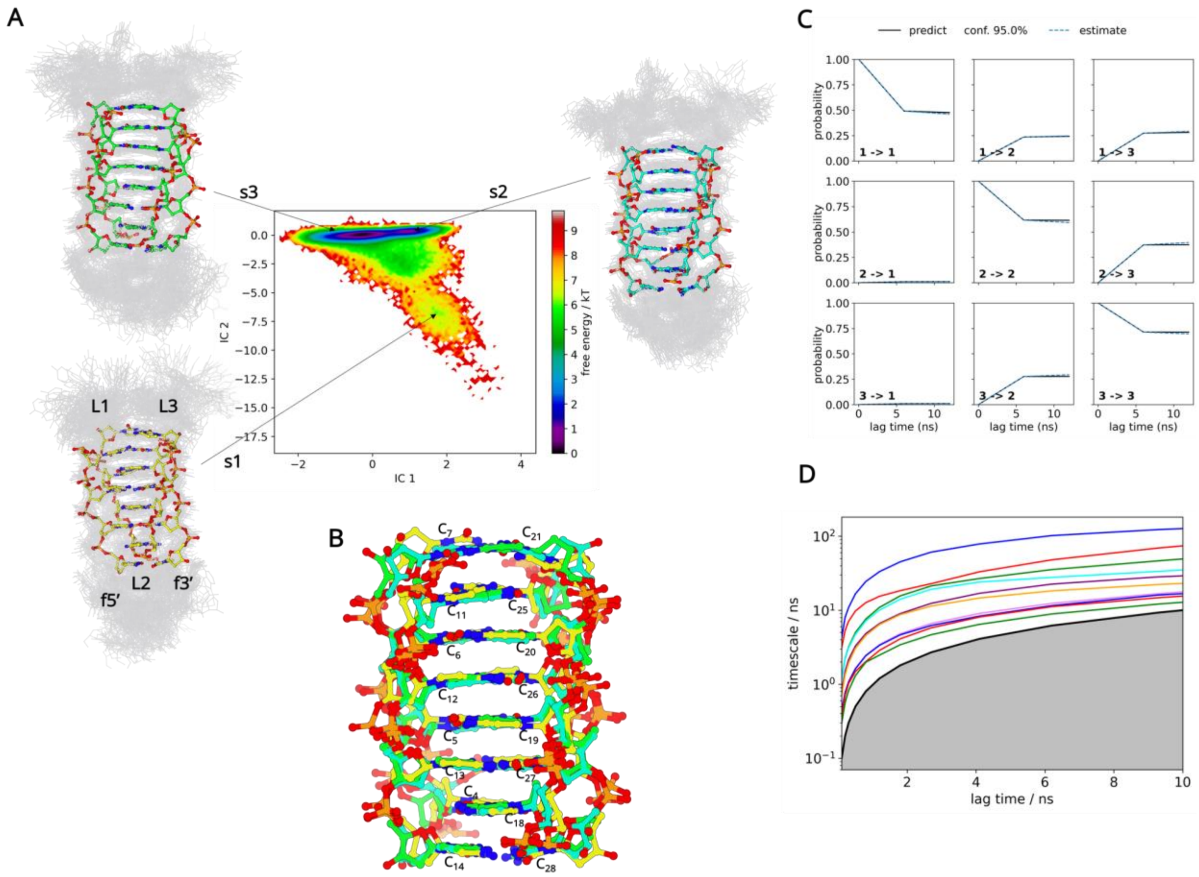


**Figure S18:** (A) Projection of the Markov state model free energy landscape along the time-lagged independent components IC1 and IC2 of 4Cdel. The representative structures s1, s2 and s3 derived from the dynamics of the central stem are marked in their basins. The positions of the loops have been marked L1, L2 and L3, while that of the 5' and 3' flanking nucleotides are labelled f5' and f3' respectively. (B) Superimposition of the three representative central stem structures of 4Cdel. (C) A Chapman-Kolmogorov (CK) plot is used to validate the self-consistency of the built MSM. 3 macrostates were defined. (D) The implied timescale (ITS) plot.

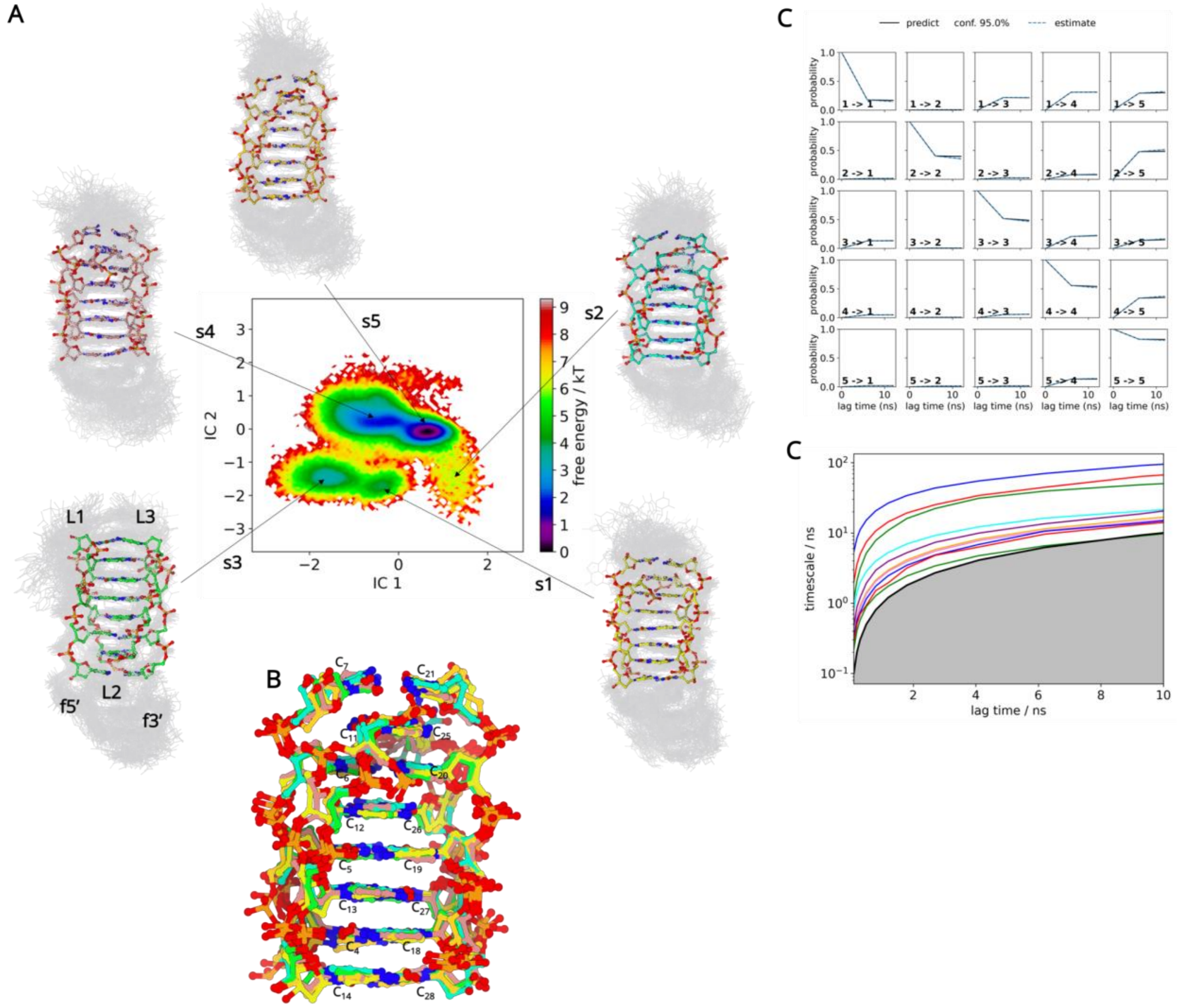


**Figure S19:** (A) Projection of the Markov state model free energy landscape along the time-lagged independent components IC1 and IC2 of 4C. The representative structures s1, s2, s3, s4 and s5 derived from the dynamics of the central stem are marked in their basins. The positions of the loops have been marked L1, L2 and L3, while that of the 5' and 3' flanking nucleotides are labelled f5' and f3' respectively. (B) Superimposition of the five representative central stem structures of 4Cdel. (C) A Chapman-Kolmogorov (CK) plot is used to validate the self-consistency of the built MSM. 5 macrostates were defined. (D) The implied timescale (ITS) plot.

### Sampling of the crystal structure conformations

The crystalline unit cell consists of two individual strands related by an inverted symmetry. The two strands (4C and 4Cdel from here onwards) are linked with the 5' end of each unit interacting with loop 3 of the other. However, the interactions in these regions are not identical; $A_2$ of strand 4C is able to form two hydrogen bonds with $A_{24}$ in strand B whereas due to a slight misalignment this interaction does not occur in the analogous region in the inverted symmetry. However, $T_1$ stabilises $A_{24}$ via a $\pi$-stacking interaction in both cases.

The ability of a simulation to fully sample its starting structure is often used as a benchmark to validate the accuracy of the sampled conformation. In our case of the ILPR i-Motif, several favourable interactions occur as a result of crystal packing and therefore the conformation found within the crystal structure may not be a realistic depiction of the structure *in vivo*. Given that once these interactions are removed, they are likely to be compensated by other interactions. Thus, the crystal structure can no longer be defined as the most stable state and may rarely be visited throughout the course of a simulation. This is particularly relevant in the case of loop 3 where crystal packing effects lead to complete solvent exposure when the structures formed in the crystalline lattice are separated. Still, assessing whether the conformations adopted by the loops, during the simulations, are able to sample their crystal structure conformation is nevertheless a useful benchmark to assess the accuracy and quality of the model created. All the loops are able to sample conformations observed in the crystalline lattice in spite of the loss of crystal packing interactions. For example, $A_8$-$A_{10}$ (loop 1), $A_{16}$-$T_{17}$ (loop 2) and $A_{22}$-$C_{21}$ (loop 3) $\pi$-stacking interactions in strand A and $C_7$-$A_8$ (loop1), $T_{15}$ $\pi$-stacking below $T_3$:$T_{17}$ base pair and $C_{21}$-$A_{22}$ (loop3) $\pi$-stacking interactions in strand B are all observed (Figure S20).

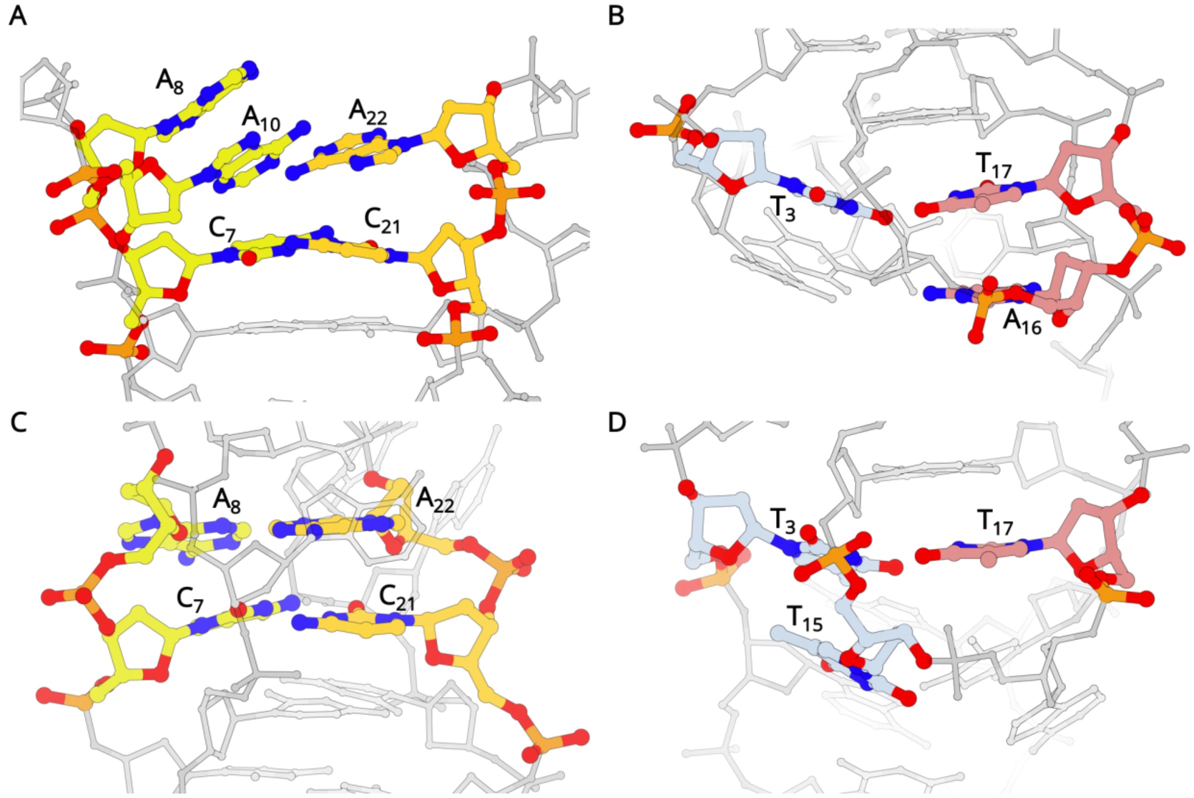


**Figure S20:** Sampling the crystallographic conformations in 4Cdel (A) Loops 1 (yellow) and 3 (orange). The non-canonical $A_{10}$:$A_{22}$ base pair caps the terminal $C_7$:$C_{21}$ base pair. $A_8$ makes $\pi$-stack with $A_{10}$; (B) Loop 2 (pink) and the flanking (cyan) nucleotides in 4Cdel. $A_{16}$ $\pi$-stacks below $T_{17}$ stabilising the $T_3$:$T_{17}$ base pair. The crystallographic conformations observed in 4C (C) The $\pi$-stacking of two non-canonical base pairs of $A_8$:$A_{22}$ above $C_7$:$C_{21}$ in Loops 1 and 3 and (D) Loop 2 and the flanking bases, where $T_{15}$ stacks below $T_3$ in 4C.

**Loop Dynamics**

Given the intrinsic rigidity of the i-motif stems enforced by the hydrogen bonding between cytosines and repulsive forces between neighbouring carbonyl and amino groups in the C·C$^+$ pair stacks, the regions either side of the stem (loop 1/3 and loop 2) can be viewed as a two different sub-systems as the dynamics in one region is likely to have a minimal effect on that of the other. Therefore, in order to assess loop conformations more directly, MSMs were created for the sequences 4C and 4Cdel in which only loop features were used as inputs for tICA, namely $\alpha/\gamma$ angles and $\chi$ angles which have proven useful previously to assess loop dynamics in G-quadruplexes.

**Loop 2**

In 4C, the most stable conformation (state 4) represents 86% of all sampled conformations. This conformation samples the interactions observed in the crystal structure including the $T_3$:$T_{17}$ base pair. Stacked below the $T_3$:$T_{17}$ base pair is the $T_5$:$A_2$:$T_{29}$ triad. In the triad, $T_{15}$ is slightly off plane. Stacking directly below $A_2$:$T_{29}$ is $T_1$:$A_{30}$ base pair. The 3' end terminal $T_{33}$ stacks below $A_{30}$. $A_{16}$ from loop 2 is solvent exposed and interacts with the 3' hydroxyl group of T33 (Figure S21). These series of base stacks between the flanking termini and loop 2 help to stabilise the structure.

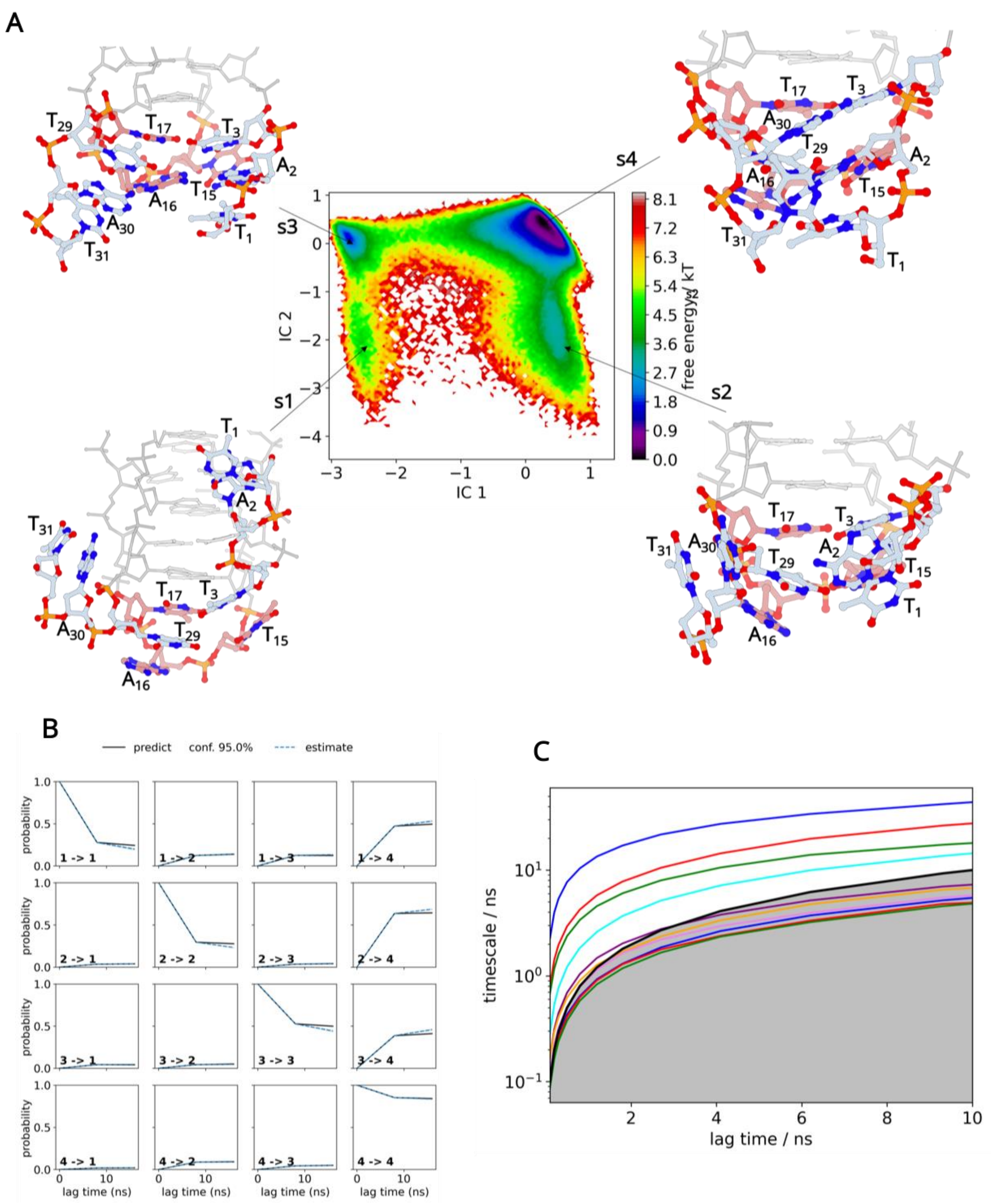


**Figure S21:** (A) The free energy landscape of Loop 2 (pink) and the 5' and 3' flanking nucleotides (cyan) in the 4C. Four structures were identified that represented their respective clusters. In the most stable conformation (s4), loop 2 interacts with the flanking region to generate a stable conformation. The extra nucleotide in the 3' flanking region enhances interactions to further stabilize the i-Motif. (B) The Chapman-Kolmogorov test and (C) the implied time scale plots.

In 4Cdel, which is missing the terminal T in the sequence, loop 2 dynamics can be seen as the interconversion between four primary states as illustrated in the free energy landscape (Figure S22).

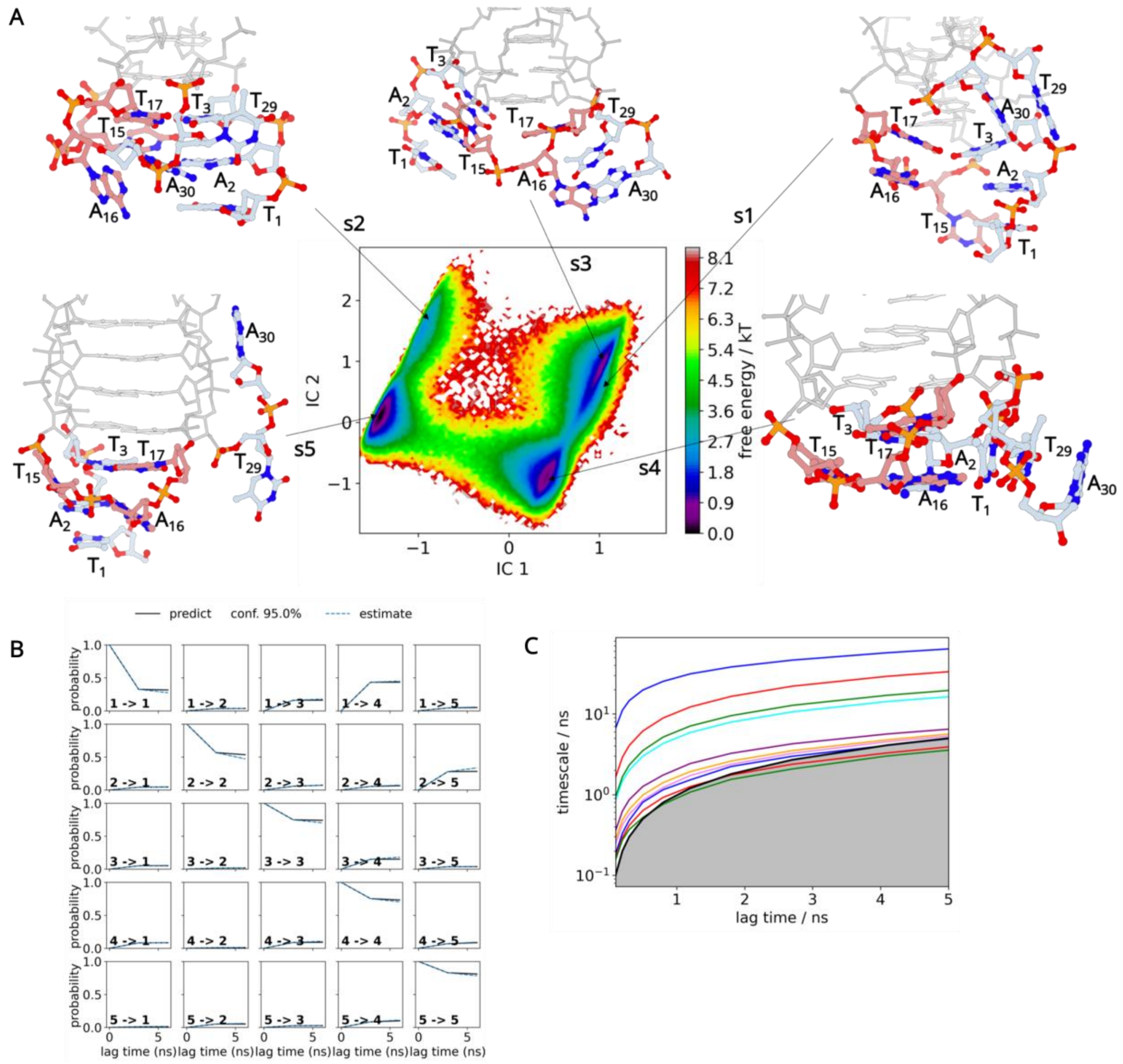

**Figure S22:** (A) The free energy landscape of Loop 2 (pink) and the 5' and 3' flanking nucleotides (cyan) in the 4Cdel. Five structures were identified that represented their respective clusters. In the most stable conformation (s4), loop 2 interacts with the flanking region to generate a stable conformation. (B) The Chapman-Kolmogorov test and (C) the implied time scale plots.

The most stable conformation (state 5), representing 39% of the equilibrium distribution system, perfectly samples the conformation displayed in the crystal structures; namely $T_{15}$ tilted inwards, $T_3$ in plane with $T_{17}$ to form a non-canonical T:T base pair, with two hydrogen bonds formed between O4 and N3 atoms that stabilises the C·C+ pair above, as well $A_{16}$ stacking below it. Interestingly $T_2$ from the 5' end makes a canonical base pair with $A_{16}$, to add further stability to the $T_3$:$T_{17}$ base pair. Finally, and as expected when crystal packing effects are removed $T_{29}$ and $A_{30}$ fold inwards and stack together (Figure S23).

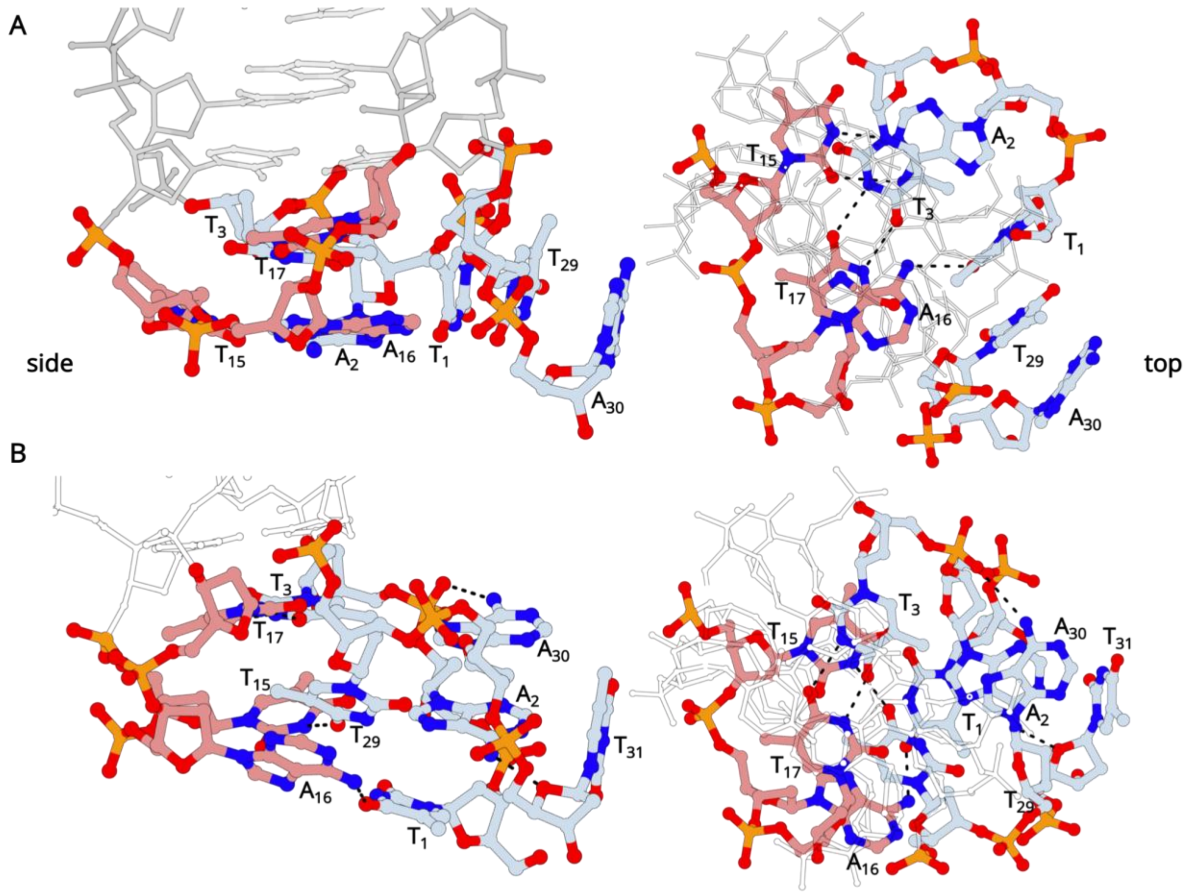


**Figure S23:** Loop2 interactions with the 5'/3' flanking nucleotides in (A) 4Cdel and (B) 4C. In 4Cdel where one nucleotide is less in the 3' flank, in addition to the $T_{13}$:$T_{17}$ base pair, $A_{16}$ makes hydrogen bond with $T_1$, $T_{15}$ with $A_2$. $T_1$ from the 5' flank is tilted inwards forming a 3-base $\pi$-stack with $T_{29}$ and $T_{30}$ from the 3' flanking nucleotides. In 4C, an extra nucleotide in the 3' flank results in enhanced hydrogen bonding and $\pi$-stacking interactions leading to a more structured capping of the end of the 4C i-motif. The side and top views if the interactions have been illustrated of Loop 2 (pink) and flanking nucleotides (cyan).

When comparing the dynamics of loop 2 between sequences 4C and 4Cdel, it becomes evident that 4C is significantly more skewed towards a single structure with metastable state 4 representing 86% of the equilibrium population. This skewing might be a result of the longer 3' end which leads to enhanced $\pi$-stacking and hydrogen bond interactions with loop 2, and thus leading to greater stability. The elongated 3' end also presents a more plausible version of what may happen *in vivo* as there are likely to be at least three residues between the start/end of the i-motif and the start of other canonical/non-canonical nucleic acid structures. It is also worth noting that the position and orientation of $T_{15}$ is unchanged across the metastable structures, highlighting to its importance in maintaining structural integrity within this region. Given its position and orientation and the interactions it makes with $A_2$, it is likely that $A_{15}$ acts like a hydrophobic cap that impedes the entrance of water into the core of the structure and therefore likely interference with the C·C$^+$ pair. Finally, our results validate the importance of non-canonical base pairs acting as capping residues in i-motif structures. All structures contain a T:T base pair between $T_3$ and $T_{17}$ that is analogous to that of the C·C$^+$ pair and thus extends the loop and further stabilised the terminal pair of the stem through favourable stacking of exocyclic carbonyls and amino groups in an antiparallel manner.

### Loops 1 and 3

To assess the dynamics of loops of 1 and 3 only, without the models being affected by the dynamics around loop 2, MSMs were then created using only the $\alpha/\gamma$ angles and $\chi$ angles of loop 1 and 3 only as they act as a singular interacting system and their dynamics should be taken together into account. The conformational space of loops 1 and 3 in strand A can be described as the interconversion between 6 metastable states (Figure S24).

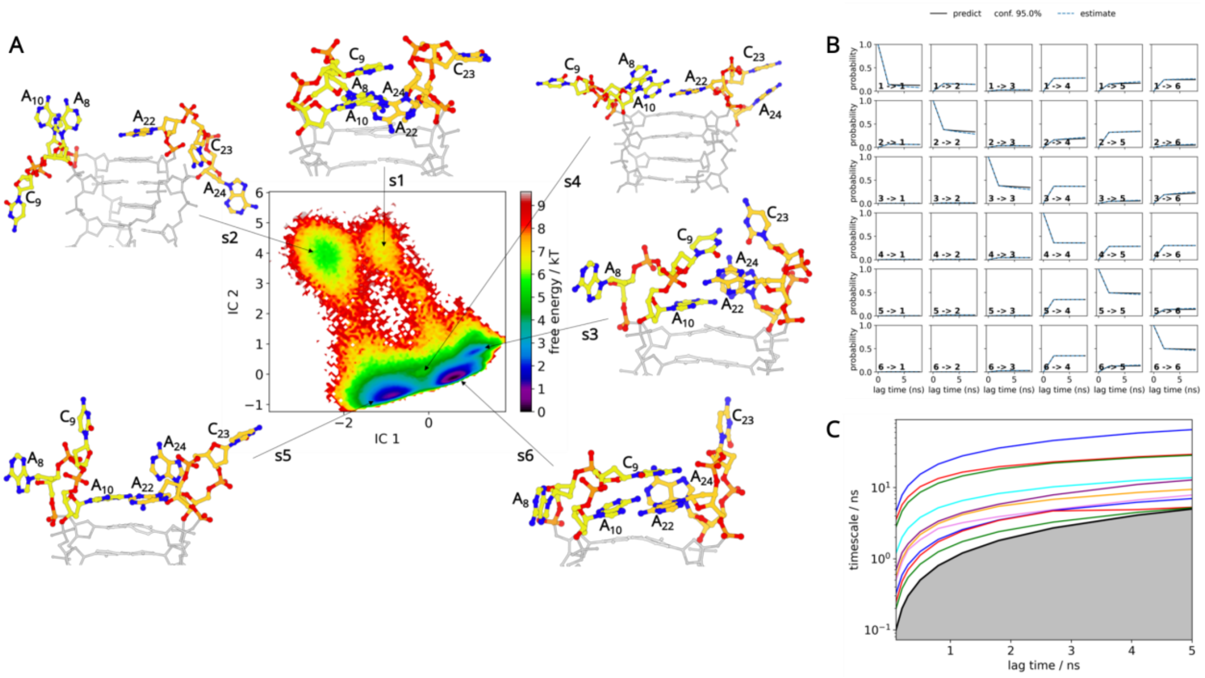


**Figure S24**: (A) The free energy landscape of Loop 1 (yellow) and Loop3 (orange) in 4Cdel structure. Six structures were identified that represented their respective clusters. Two of these (s5 and s6) correspond to low energy conformation. In the most stable conformation (s6), C9 from Loop 1 stacks on top of $A_{22}$ to stabilise the $A_{10}$:$A_{22}$ non-canonical base pair. (B) The Chapman-Kolmogorov test and (C) the implied time scale plots.

The most populated state is state 6 accounting for 52% of all sampled conformations. In loop 1, $A_{10}$ is able to stack on $C_7$ to form a non-canonical base pair interaction with $A_{22}$, with hydrogen bonds formed between the N6 and N7 atoms in both bases. Interestingly, in this cluster we observe stacking of $C_9$ above $A_{22}$ and $A_8$ above $A_{10}$. Metastable state 6 recapitulates the conformations and interaction that are observed in the crystal structure (Figure S20). State 3 is a substate of state 6, where $C_9$ stacks above $A_{10}$:$A_{22}$ base pair. In metastable state 5 (38% population), the $A_{10}$:$A_{22}$ base pair is formed but there is no stacking observed by $A_8$ or $C_9$ bases above the $A_{10}$:$A_{22}$ base pair. State 4 is a substate of metastable state 5, where the $A_{10}$:$A_{22}$ base pair is not formed, while $A_8$ makes $\pi$-stacking with $A_{10}$. State 1 is a low populated, high energy conformation where loop 1 and 3 are collapsed on the terminal $C_7$:$C_{21}$ base pair. The $A_{10}$:$A_{22}$ base pair that stacks on top of $C_7$:$C_{21}$ base pair is off centred, while $C_9$ stacks on top of $A_{10}$:$A_{22}$ base pair instead of A8, which is tucked in a groove and is shielded from the solvent by $A_{24}$. Nucleotide $A_{23}$ is solvent exposed. State 2 population represents the open conformation of the loop 1 and 3. In this state, the loops move outward exposing the $A_{10}$:$A_{22}$ base pair and in some instances the $C_7$:$C_{21}$ base pairs.

Loop 1 and 3 in 4C are separated by tICA into 4 metastable states (Figure S25). State 1 in 4C represents the crystalline conformation where $A_8$:$A_{22}$ base pair is present, while bases $C_9$, $A_{10}$, $C_{23}$ and $A_{24}$ are solvent exposed. In state 2, the $A_8$ and $A_{22}$ base pair is lost and the sugar phosphate backbone of loops 1 and 3 is in a constrained conformation pointing towards the central i-Motif stem. Unlike 4Cdel, loop 1 in the most populated state 3 (91%) of 4C does not fully sample the conformation found in the crystal structure. The $A_8$:$A_{22}$ base pair is abolished and $A_{22}$ is instead stabilised by forming two hydrogen bonds with its N6 amine, making interactions with the O4' and N3 atoms in $A_{10}$. $A_8$ is stabilised through the contact between its N6 amine and the phosphate oxygen in $A_{24}$. $A_{23}$ and $A_{24}$ stack next to each other in a vertical orientation, this stacking arrangement is further stabilised by the interaction between the $A_{24}$ N6 amine and the $C_{23}$ phosphate group. Furthermore, the slight inward rotation of loop 3 allows the stabilisation of the $C_7$ N4 amine via interaction with the $A_{24}$ phosphate. State 4 is similar to state 3. The only difference is the arrangement of $C_9$, $A_{10}$, $C_{23}$ and $A_{24}$ bases in loops 1 and 3 respectively.

To summarise, upon inspection of structures extracted from coarse-grained models, 4Cdel featured far more unstructured conformations in loops 1 and 3 while those in 4C, with the full flanking sequence, are well ordered.

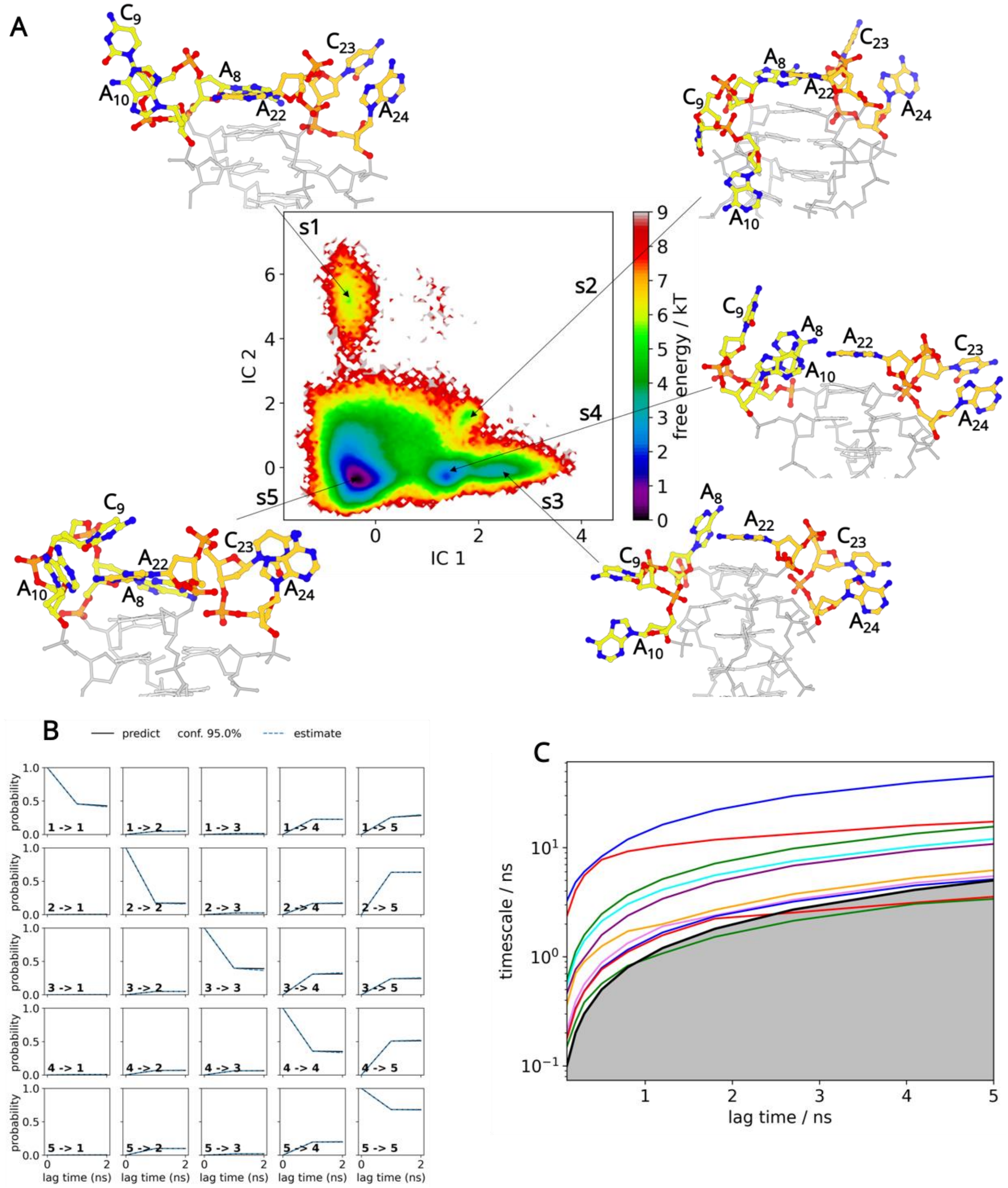


**Figure S25**: (A) The free energy landscape of Loop 1 (yellow) and Loop3 (orange) in the 4C structure. Five structures were identified that represented their respective clusters. s5 represents the most stable state where A8:A22 base pair is formed while $C_9$ from loop 1 makes π-stacking interactions with $A_8$. Loop 3 is stabilised by π-stacking interactions between $C_{23}$ and $A_{24}$. (B) The Chapman-Kolmogorov test and (C) the implied time scale plots.
